# HiC-LEGO: Biologically Guided High-Resolution 3D Genome Reconstruction Preserves Chromatin Organization at Kilobase Resolution

**DOI:** 10.64898/2026.08.30.748130

**Authors:** Abhishek Pandeya, Md Fahimul Kabir Chowdhury, Oluwatosin Oluwadare

## Abstract

Three-dimensional (3D) chromosome reconstruction from Hi-C contact maps remains challenging because genome organization is hierarchical and fine-resolution models must reconcile local structure with chromosome-scale constraints. Here we present HiC-LEGO, a domain-aware hierarchical framework integrating ensemble chromatin domains with graph-based structural learning and progressive chromosome assembly. By combining ensemble domain selection with hierarchical reconstruction, HiC-LEGO reduces dependence on individual domain definitions while maintaining local organization during chromosome-scale assembly. Across five human cell lines at 5-kb resolution, HiC-LEGO achieves higher reconstruction concordance than evaluated state-of-the-art methods while better preserving domain organization. At 1-kb resolution, HiC-LEGO reconstructs complete GM12878 chromosome 8 and recovers close spatial proximity between an epigenomically supported distal MYC enhancer and its promoter. Reconstructions from 5-kb Micro-C data show that 249 experimentally defined RCMC microcompartment interactions at the Ppm1g locus occupy compact 3D configurations. In the breast cancer dataset, HiC-LEGO reconstructs structures that maintain stable TAD organization across healthy breast, primary tumors, and liver metastases, while revealing greater inter-patient structural heterogeneity in malignant pleural effusion samples. Pore-C validation shows that experimentally observed multi-way contacts spanning 1–5 Mb are enriched in compact reconstructed configurations across all 23 chromosomes. Thus, HiC-LEGO preserves regulatory interactions, disease-associated chromatin organization and higher-order spatial relationships beyond pairwise contact-map concordance.

## 1 Introduction

The three-dimensional (3D) organization of chromosomes plays a fundamental role in gene regulation, genome stability, and other cellular processes [1]. Topologically associating domains (TADs) represent an important level of this organization, defining genomic regions with enriched internal interactions and contributing to regulatory insulation [2]. To characterize these spatial interactions, chromosome conformation capture (3C)-based methods have been developed, among which Hi-C enables genome-wide mapping of chromatin contacts across genomic scales [1–3]. These contact maps provide a basis for reconstructing 3D chromosome structures, although translating interaction frequencies into accurate spatial coordinates remains computationally challenging. Numerous reconstruction methods have been developed, initially focusing largely on coarser genomic resolutions (typically *≥*25 kb), and have been organized into different methodological classes according to how contact information is translated into 3D structure [4–6].

Oluwadare et al. [6] categorized 3D genome reconstruction algorithms into three primary methodological classes: distance-based, contact-based, and probability-based methods. Distance-based approaches convert interaction frequencies (IFs) into spatial distances and optimize genomic coordinates accordingly. LorDG [7] uses a nonlinear Lorentzian objective, 3DMax [8] applies maximum-likelihood optimization, ChromSDE [9] uses semidefinite programming, and ShNeigh [10] combines multidimensional scaling with spatial dependence between neighboring loci. Contact-based methods, such as MOGEN [11] and Gen3D [12], avoid explicit IF-to-distance conversion and instead optimize structures to satisfy contact-derived constraints. Probability-based methods model interaction counts statistically; PASTIS [13] uses a Poisson likelihood, whereas BACH [14] applies Bayesian inference to estimate chromosome conformations.

MacKay and Kusalik [15] described reconstruction methods from a algorithmic perspective, grouping the methods into dimensionality reduction, graph/network theory, maximum-likelihood estimation, and statistical modeling. In this classification, dimensionality-reduction approaches project high-dimensional contact relationships into 3D space, graph-based approaches represent genomic loci and their interactions as nodes and edges, maximum-likelihood methods optimize structures according to a likelihood function, and statistical approaches infer conformations using probabilistic models. Individual reconstruction tools may combine strategies from more than one of these categories [15]. More recent approaches integrate experimental or biophysical information: GEM-FISH [16] combines Hi-C and FISH measurements with polymer constraints for hierarchical chromosome reconstruction, whereas Hi-BDiSCO [17] incorporates Hi-C or Micro-C restraints into Brownian-dynamics simulations for high-resolution mesoscale chromatin modeling.

For high-resolution (*<*25 kb) 3D chromosome and genome structure reconstruction, relatively few methods have been developed. To our knowledge, two algorithms have specifically targeted chromosome-scale reconstruction at these resolutions: Hierarchical3DGenome [18], which reconstructs TAD-level structures and assembles them into complete chromosomes using inter-TAD contacts, and FLAMINGO [19], which enables high-resolution reconstruction through modular modeling and optimization-based embedding. High-resolution reconstruction is challenging because finer binning substantially increases the number of modeled loci and structural constraints while requiring the reconstructed coordinates to preserve finescale features such as chromatin loops different range of interactions. To manage this complexity, recent approaches have adopted hierarchical strategies that reconstruct smaller regions before assembling them into chromosome-scale structures [18, 19]. However, this makes the definition of those regions a critical part of the reconstruction process. Existing hierarchical approaches address domain definition in different ways, but both can limit biological fidelity. Some approaches partition chromosomes using fixed genomic intervals or construct a coarse backbone before local refinement, which can split or merge biologically defined TADs [19]. Others rely on domains identified by a single TAD caller [18], allowing caller-specific boundary errors to propagate into the reconstructed structure. This is important because TAD-detection methods use substantially different assumptions, including insulation score, statistical modeling, clustering, and graph-based modularity approaches, and can consequently produce different domain boundaries [20–22]. These limitations motivate high-resolution reconstruction strategies that preserve biologically meaningful domain organization while remaining robust to uncertainty in TAD detection.

To address these limitations, we introduce HiC-LEGO, a biologically guided, TAD-aware framework for high-resolution (*<*25 kb) 3D genome reconstruction. HiC-LEGO combines graph neural network-based structural learning with ensemble domain selection and hierarchical chromosome assembly. HiC-LEGO integrates outputs from multiple TAD-detection methods to identify consensus domains, reducing dependence on individual domain definitions and limiting the propagation of caller-specific boundary errors. Graphbased learning captures structural relationships among interacting chromatin regions, while the selected domains guide reconstruction and progressive assembly from local structures to complete chromosomes. These components directly address the domain-definition and local-to-global assembly limitations of existing high-resolution approaches while providing a biologically grounded framework for chromosome-scale reconstruction.

## 2 Results

### 2.1 HiC-LEGO for High Resolution Reconstruction of 3D Genome structures

HiC-LEGO is a unified framework for high-resolution 3D chromosome reconstruction that combines ensemble domain selection, graph-based modeling, and hierarchical assembly. It uses a graph-based model combining mean-neighbor aggregation with LINE-derived structural embeddings to infer chromatin representations. These learned representations form the basis of the subsequent TAD-guided fine-resolution reconstruction.

As shown in Fig. 1a–c, HiC-LEGO first identifies an optimal set of topologically associating domains (TADs) by integrating calls from multiple TAD-detection methods and comparing them using the Measure of Concordance framework [23] (See section 4.1). HiC-LEGO then reconstructs high-resolution 3D structures for the selected domains using learned graph embeddings and hierarchically assembles them, first within megabase-scale LEGOs and then into a chromosome-wide structure anchored by a 1-Mb backbone. The visualization of the 3D structure was done using Chimera [24].

**Fig 1.**
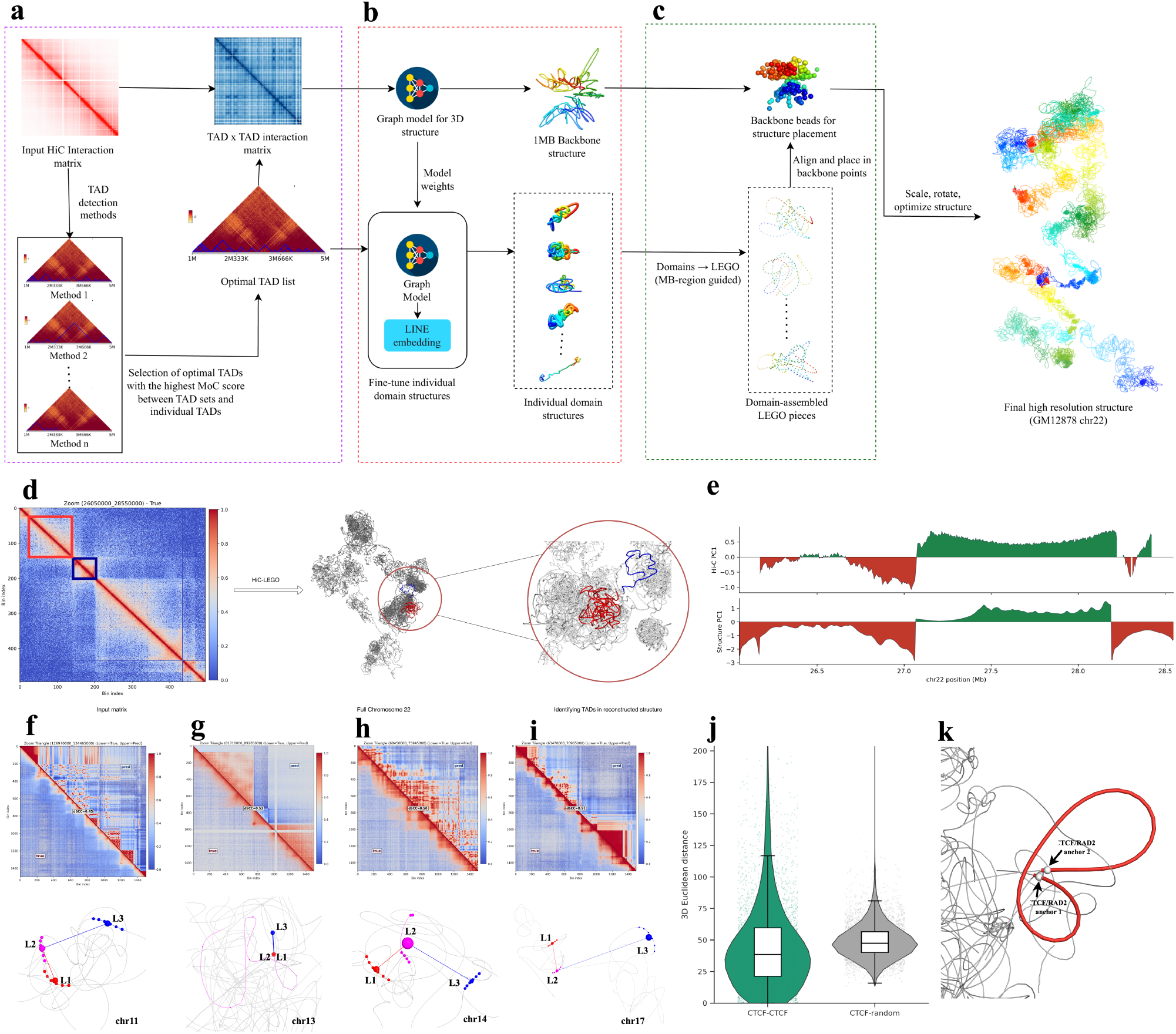
HiC-LEGO framework and structural validation of reconstructed chromosomes. a,b,c. HiC-LEGO pipeline overview. d. Representative 2.5-Mb region of chromosome 22 (26.05–28.55 Mb), where prominent contact domains in the input Hi-C map correspond to spatially localized regions in the reconstructed chromosome structure. e. Hi-C-and structure-derived PC1 profiles at the same GM12878 chr22 locus, showing preservation of compartment-scale organization in the reconstructed structure. f-i. GM12878 FISH loci showing the corresponding genomic regions and the reconstructed 3D structures, with measured distances between the L1–L2 and L2–L3 probe pairs. j. Distribution of 3D distances between CTCF–CTCF loop anchors and distance-matched random pairs. k. Representative reconstructed CTCF/RAD21-mediated loop with labeled anchor regions.

Local domain organization remained identifiable after chromosome-scale assembly. In chromosome 22, a 2.5-Mb region at 26.05–28.55 Mb contained prominent contact domains in the input Hi-C map that corresponded to spatially localized regions in the reconstructed chromosome (Fig. 1d), visualized usin PyMOL [25]. Their relative organization was retained within the full chromosome structure, which was used for locus-level validation. At 5-kb resolution, PC1 compartment profiles computed with cooltools [26] showed that HiC-LEGO preserved broad compartment organization in GM12878 chromosome 22 after chromosome-scale assembly (Fig. 1e).

We then evaluated experimentally characterized local spatial relationships. Representative 7.5-Mb GM12878 regions showed agreement between observed and reconstructed contact organization, with regional distance Spearman correlation coefficient (dSCC) values of 0.45–0.53 (Fig. 1f–i). These regions corresponded to FISH-characterized loci on chromosomes 11, 13, 14, and 17 from Rao et al. [27]. Mapping the L1, L2, and L3 probe regions onto the reconstructed structures reproduced the expected ordering in which the loop-associated L1–L2 pair was spatially closer than L2–L3 (Fig. 1f–i; Supplementary Table S1), providing orthogonal validation beyond contact-map concordance.

We found the chromatin loops which are strongly associated with CTCF and cohesin components such as RAD21, with loop-anchor CTCF motifs predominantly arranged in convergent orientation [1, 28]. In the reconstructed structures, CTCF–CTCF anchor pairs showed shorter 3D distances than genomic-distance-matched random pairs (Fig. 1j; Supplementary Table S2). A representative CTCF/RAD21-supported loop placed the two annotated anchors in close spatial proximity, connected by a compact chromatin path (Fig. 1k; Supplementary Table S2). Thus, these analyses show that HiC-LEGO preserves structural organization from chromosome-scale domains to experimentally characterized local chromatin contacts.

### 2.2 Ensemble-based TAD optimization establishes a robust consensus domain framework for 3D genome reconstruction

To evaluate the reconstruction accuracy of HiC-Lego at 5 kb resolution, we first optimized the TAD selection strategy used in the reconstruction pipeline. Since TADs provide the structural units used to guide local chromatin folding, the choice of domain set can directly influence the final 3D model. We therefore compared the reconstruction performance obtained using the proposed ensemble-based domain selection hypotheses (See section 4.1). We excluded Hypothesis 1 from this quantitative comparison because the strict requirement that domains be supported by all TAD-calling methods produced very few retained TADs, which was insufficient genomic coverage for reliable 3D reconstruction. We then compared Hypothesis 2 and Hypothesis 3 across GM12878 chromosomes using dSCC, which measures the agreement between the observed contact matrix and the contact matrix inferred from reconstructed 3D distances.

Across all 22 autosomes and chromosome X, the reconstruction represented by Hypothesis 2 consistently achieved higher distance Spearman correlation coefficients (dSCCs) than Hypothesis 3 (See Supplementary Figure S1). The mean chromosome-wide dSCC increased from 0.399 under Hypothesis 3 to 0.477 under Hypothesis 2, with the same direction of improvement observed for every chromosome examined. This consistent gain across chromosomes, rather than an improvement driven by a small subset of loci, supported the use of the Hypothesis 2 assembly strategy for subsequent HiC-LEGO reconstruction.

### 2.3 Benchmarking HiC-LEGO Against State-of-the-Art High-Resolution 3D Genome Reconstruction Methods at 5 kb resolution across cell lines

We compared HiC-LEGO with FLAMINGO [19] and Hierarchical3DGenome (H3DG) [18] at 5-kb resolution across GM12878, K562, HMEC, IMR90, and HUVEC. For each cell line, all 22 autosomes and chromosome X were reconstructed and evaluated, yielding 23 chromosomes per cell line and 115 chromosome-level reconstructions in the pooled analysis. We measured reconstruction accuracy using the distance Spearman correlation coefficient (dSCC), which quantifies agreement between distances implied by the input contact map and those recovered from the reconstructed 3D structure. Across the 115 pooled chromosome-level values, HiC-LEGO showed the highest dSCC distribution and a reduced low-accuracy tail relative to H3DG (Fig. 2a).

**Fig 2.**
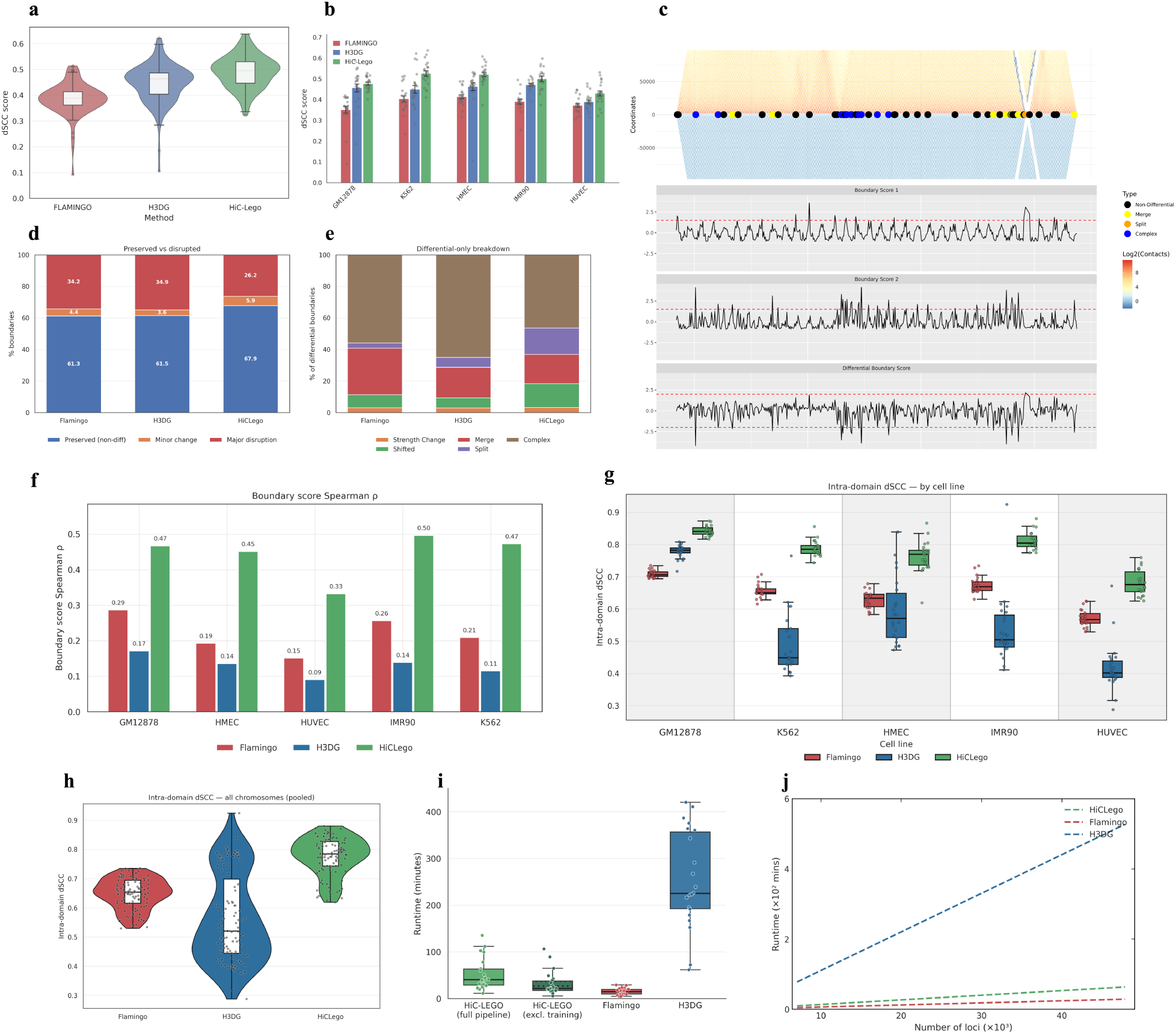
HiC-LEGO improves reconstruction accuracy and preserves domain organization. (a) Genome-wide dSCC distributions showing higher reconstruction concordance for HiC-LEGO than FLAMINGO and H3DG. (b) Per-cell-line dSCC, showing consistently higher HiC-LEGO accuracy across all five cell lines (bars *±* SD; points = chromosomes). (c) GM12878 chr22:26.05–28.55 Mb example showing correspondence between contact domains in the input and reconstructed maps, together with boundary classes and boundary-score profiles. (d) HiC-LEGO preserves the largest fraction of non-differential boundaries while reducing major disruptions. (e) Composition of differential boundaries classified as strength change, shifted, merge, split, or complex. (f) Higher boundary-score Spearman *ρ* for HiC-LEGO across all five cell lines indicates improved preservation of local domain organization. (g) Consistently higher intra-domain dSCC across individual cell lines. (h) Pooled intra-domain dSCC showing higher within-domain reconstruction accuracy for HiC-LEGO. (i) Per-chromosome runtime comparison, including HiC-LEGO with and without training time. (j) Runtime scaling with increasing numbers of reconstructed loci for HiC-LEGO, FLAMINGO, and H3DG.

We observed the same trend within each cell line (Fig. 2b). HiC-LEGO achieved the highest mean chromosome-level dSCC across all 23 chromosomes in each of the five cell lines. The largest gains were observed in K562 and HMEC, where HiC-LEGO reached 0.52, compared with 0.44–0.45 for H3DG and 0.40–0.41 for FLAMINGO. HiC-LEGO improved mean dSCC in GM12878 and IMR90, with a smaller but consistent gain in HUVEC. Chromosome-level values showed that these improvements were distributed across the evaluated chromosomes rather than driven by only a few cases. We next compared computational runtime across the three methods. Despite its graph-based learning framework, HiC-LEGO remained computationally competitive with FLAMINGO, an optimization-based low-rank matrix-completion method [19], while being substantially faster than H3DG (Fig. 2i). HiC-LEGO is also GPU-compatible, providing additional scalability for larger and higher-resolution reconstructions. Overall, it combines consistently higher 5-kb reconstruction accuracy with practical chromosome-scale computational performance.

### 2.4 HiC-LEGO preserves TAD boundary architecture and intra-domain organization

Since HiC-LEGO uses topologically associating domains (TADs) as structural units during chromosome assembly [2], used TADCompare [29] to assess boundary agreement, which classifies boundaries as preserved or altered through shifts, strength changes, merges, splits, and complex rearrangements. A representative GM12878 region shows concordant boundary calls and boundary-score profiles between the input and reconstructed maps (Fig. 2c).

HiC-LEGO preserved a larger fraction of TAD boundaries than either comparison method (Fig. 2d). Overall, 67.9% of HiC-LEGO boundaries were non-differential, compared with 61.3% for FLAMINGO and 61.5% for H3DG. Major disruptions occurred at 26.2% of HiC-LEGO boundaries, versus 34.2% and 34.9% for FLAMINGO and H3DG, respectively. Among differential boundaries, complex changes were the most common, while shifts, merges, splits, and strength changes occurred less frequently (Fig. 2e). These results indicate that HiC-LEGO more often preserves the genomic positions separating neighboring chromatin domains and less frequently introduces large changes in domain partitioning.

Boundary-strength profiles were better preserved by HiC-LEGO (Fig. 2f). Spearman correlations between input and reconstructed boundary scores were 0.47 in GM12878, 0.45 in HMEC, 0.33 in HUVEC, 0.50 in IMR90, and 0.47 in K562. In comparison, FLAMINGO produced correlations of 0.15–0.29 and H3DG correlations of 0.09–0.17. Thus, HiC-LEGO retained not only boundary positions, but the relative strength with which neighboring domains remained separated in the reconstructed contact organization.

Pooled intra-domain dSCC values were highest for HiC-LEGO (Fig. 2g), and we observed it independently in GM12878, K562, HMEC, IMR90, and HUVEC (Fig. 2h). Higher intra-domain dSCC indicates that the internal contact organization of individual TADs is more faithfully retained after chromosome-scale assembly. These analyses show that HiC-LEGO preserves both the boundaries separating chromatin domains and the interaction structure within them.

### 2.5 HiC-LEGO resolves regulatory chromatin proximity at 1 kb resolution

We reconstructed the complete GM12878 chromosome 8 at 1 kb and evaluated the MYC locus using HiC-LEGO. Although chromosome-scale 1-kb reconstruction has been demonstrated previously [19], biological validation at this resolution remains limited. We assessed both chromosome-wide contact concordance and spatial recovery of an epigenomically supported regulatory interaction in 1kb structure built using HiC-LEGO.

The MYC region contains the promoter, distal enhancer-associated chromatin, and long-range loops identified in high-resolution GM12878 Hi-C data [1]. We selected a HiCCUPS loop spanning 525 kb between chr8 *≈* 128.220–128.225 Mb and chr8 *≈* 128.745–128.750 Mb (Fig. 3a,b). The distal anchor overlaps chr8 *≈* 128.216–128.230 Mb, supported by a ChromHMM strong-enhancer state [30] and GM12878 H3K27ac enrichment [31].

**Fig 3.**
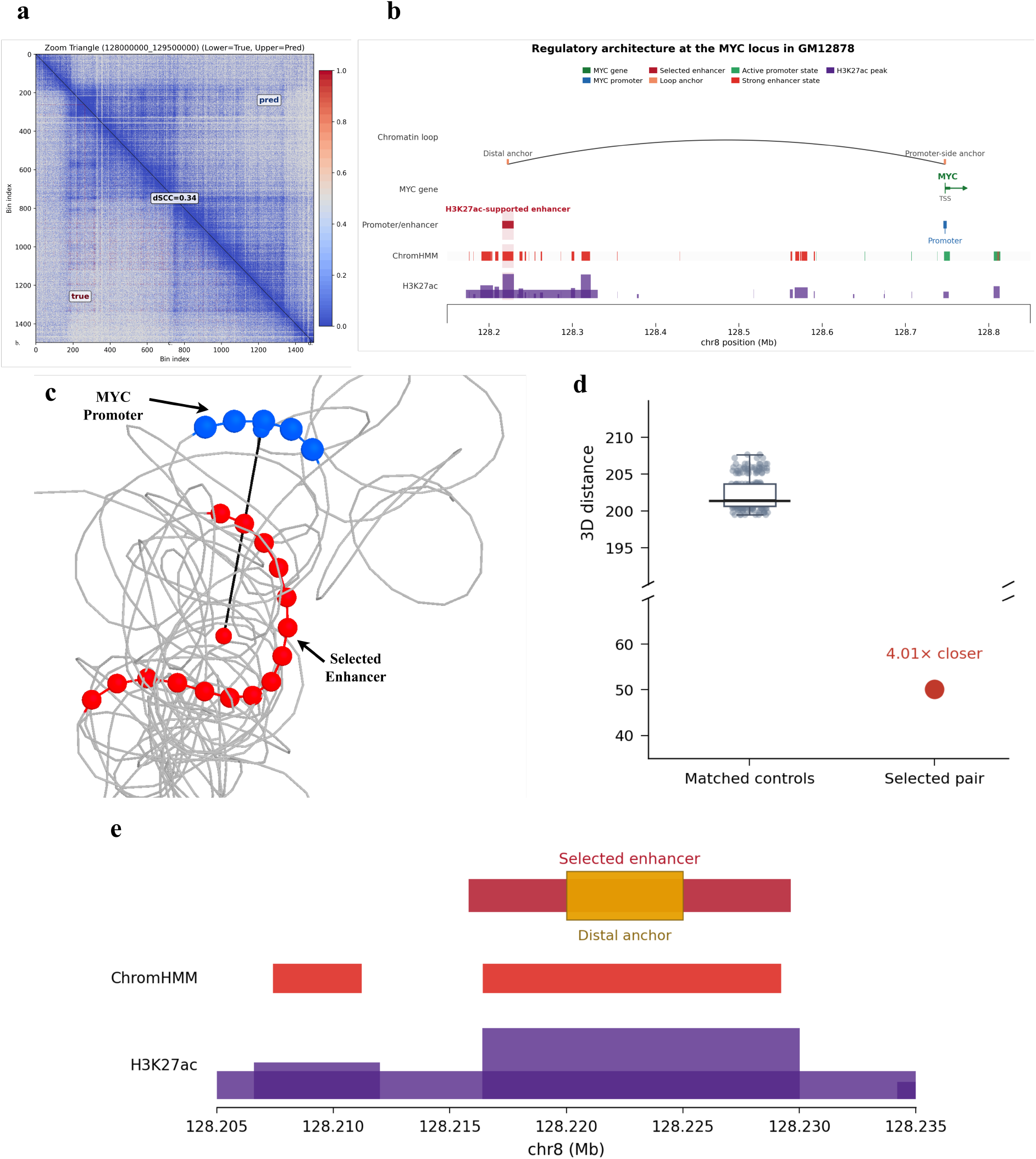
HiC-LEGO recovers regulatory organization at the MYC locus in the GM12878 chr8 1-kb reconstruction. (a) Observed Hi-C contacts (lower triangle) and structure-derived contacts (upper triangle) across chr8:128.0–129.5 Mb, with chromosome-wide dSCC = 0.34. (b) Regulatory architecture of the MYC locus, showing a HiCCUPS loop spanning ~525 kb between a distal anchor (chr8:128.220–128.225 Mb) overlapping an H3K27ac-supported strong-enhancer region and a promoter-side anchor (chr8:128.745– 128.750 Mb) near MYC. (c) Reconstructed 3D orga8nization places the selected enhancer (red) in spatial proximity to the MYC promoter (blue). (d) The promoter–enhancer pair has a 3D distance of 50.17, 4.01 *×* smaller than the median of 200 genomic-distance-matched controls (201.39; empirical *p* = 0.0050). (e) Enlarged view (visualized using PyMOL [25]) of the distal region showing overlap of the selected enhancer and loop anchor with ChromHMM strong-enhancer states and H3K27ac signal.

The 1-kb reconstruction retained chromosome-wide and local contact organization, with a chromosome 8 dSCC of 0.34 (Fig. 3a). Although lower than at 5 kb, this shows that measurable contact structure remains recoverable after fivefold finer genomic binning and a corresponding increase in modeled loci. The MYC region retained local diagonal enrichment and interaction patterns present in the observed map.

The selected enhancer–promoter pair was separated by a 3D Euclidean distance of 50.17, compared with a median of 201.39 across 200 genomic-distance-matched controls (Fig. 3c,d). The pair was 4.01-fold closer than the control median and lay at the extreme compact end of the distribution (empirical *p* = 0.0050). This spatial proximity is consistent with the long-range regulatory interaction identified in experimental Hi-C data [1].

The distal region was independently supported by a ChromHMM strong-enhancer state [30] and prominent GM12878 H3K27ac enrichment [31] (Fig. 3e). Thus, the compact 3D interaction connects the MYC promoter to an epigenomically supported distal regulatory region rather than to an arbitrary genomic segment.

These results show that HiC-LEGO can reconstruct a complete human chromosome at 1-kb resolution while retaining measurable chromosome-scale contact organization and biologically meaningful regulatory proximity. We limited this analysis to chromosome 8 because of the higher computational cost of chromosome-wide modeling at 1 kb.

### 2.6 HiC-LEGO captures Micro-C-resolved chromatin architecture and compact RCMC microcompartment interactions at Ppm1g locus

We used HiC-LEGO on mouse embryonic stem-cell (mESC) Micro-C data in order to utilize local interaction information provided by Micro-C to generate a structure. Micro-C uses micrococcal nuclease digestion and resolves chromatin contacts at finer genomic scales than conventional restriction enzyme-based Hi-C [32]. We reconstructed structures independently from mESC Hi-C and Micro-C maps at 5-kb resolution. At the Sox2 locus on chromosome 3, Micro-C showed sharper short-range contact blocks and focal interactions than Hi-C, and these features were retained in the contact map derived from the Micro-C reconstruction (Fig. 4a).

**Fig 4.**
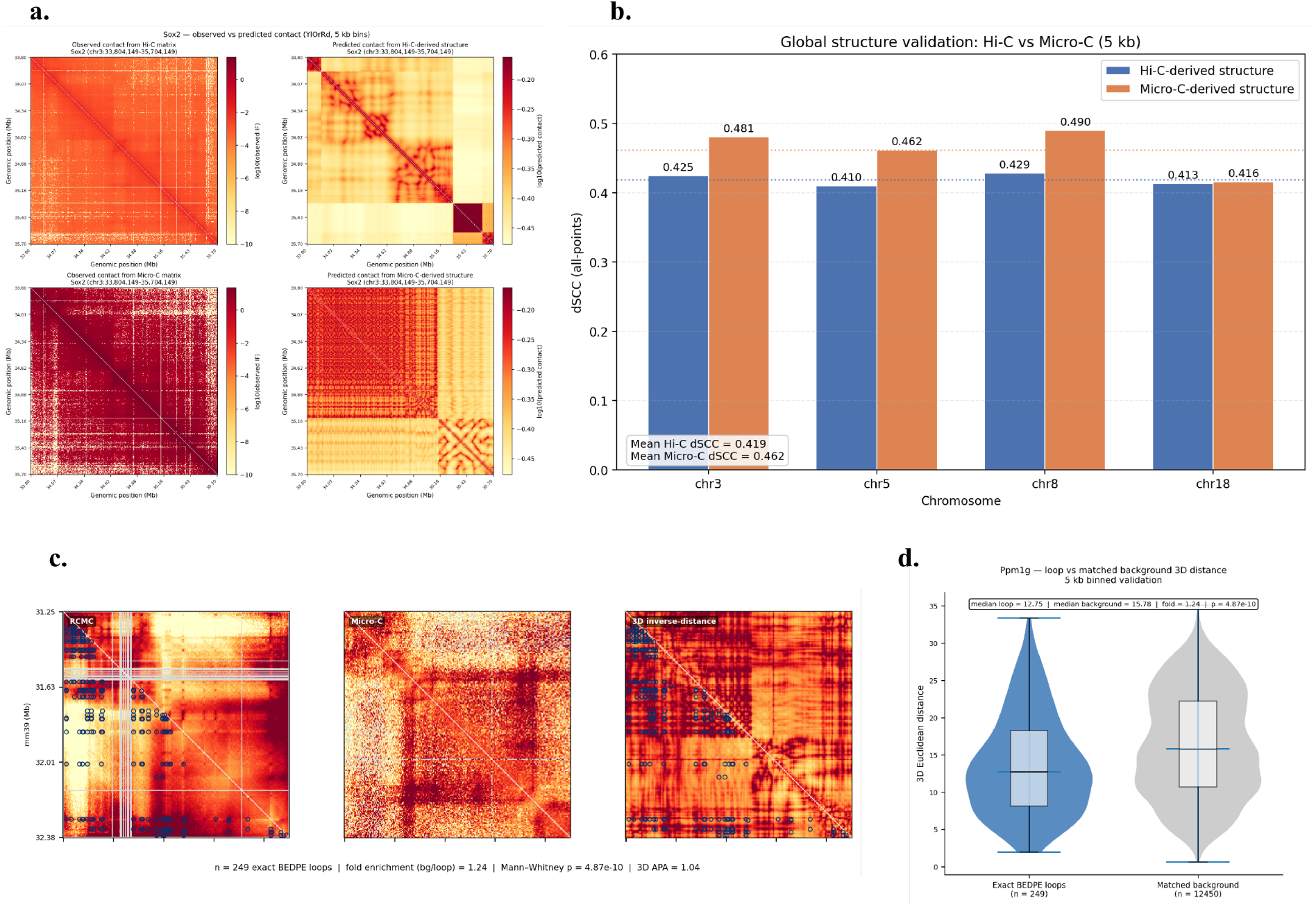
HiC-LEGO captures Micro-C–resolved fine-scale chromatin organization and preserves microcompartment interactions. (a) Sox2 locus (chr3 *≈* 33.80–35.70 Mb), showing observed and structure-derived contact maps from Hi-C (top) and Micro-C (bottom); the Micro-C reconstruction captures finer local contact patterns. (b) Full-chromosome dSCC at 5 kb for chr3, chr5, chr8, and chr18, with higher mean concordance for Micro-C-derived than Hi-C-derived structures (0.462 vs 0.419). (c) Ppm1g locus (chr5 *≈* 31.25–32.38 Mb), showing correspondence among experimentally defined RCMC microcompartment interactions (*n* = 249), Micro-C contacts, and structure-derived inverse-distance contacts; 3D APA = 1.04. (d) RCMC-defined microcompartment pairs are more spatially compact than genomic-distance-matched background pairs (median 3D distance 12.75 vs 15.78; 1.24-fold enrichment; one-sided Mann–Whitney *p* = 4.87 *×* 10^−10^).

At the chromosome scale, Micro-C-derived structures achieved a higher mean all-points dSCC than Hi-C-derived structures across chromosomes 3, 5, 8, and 18, corresponding to the chromosomes containing the RCMC capture loci analyzed here (0.462 versus 0.419; Fig. 4b). dSCC increased from 0.425 to 0.481 on chromosome 3, 0.410 to 0.462 on chromosome 5, and 0.429 to 0.490 on chromosome 8, while chromosome 18 remained comparable (0.413 versus 0.416). We used the corresponding mESC Hi-C data from Bonev et al. [33] for comparison. These results show that HiC-LEGO can leverage the finer-scale contact information provided by Micro-C to improve chromosome-scale 3D reconstruction concordance, consistent with the enhanced local chromatin resolution reported for Micro-C [32].

We examined the Ppm1g locus using published Region Capture Micro-C (RCMC) microcompartment annotations [34]. We analyzed 249 exact BEDPE-defined interaction pairs across 1.1 Mb (Fig. 4c). These interactions aligned with focal features in the RCMC and Micro-C maps and were visible in the inverse-distance contact representation of the reconstructed structure.

The 249 annotated interactions were spatially closer in the reconstruction than 12,450 genomic-distance-matched background pairs (see Supplementary Methods), with median 3D distances of 12.75 and 15.78, respectively. This corresponds to a 1.24-fold enrichment in spatial proximity (one-sided Mann–Whitney U test, *p* = 4.87 *×* 10^−10^; Fig. 4d). The 3D aggregate peak analysis showed a smaller enrichment (APA = 1.04), indicating that the main signal is a broad shift toward shorter interaction-anchor distances rather than a strong focal aggregate peak. Our analysis shows that experimentally defined microcompartment interactions are spatially compact in the reconstructed structures, with RCMC-defined contacts [34] forming more compact 3D configurations than matched controls. These results demonstrate that HiC-LEGO preserves fine-scale chromatin organization in Micro-C-derived 3D reconstructions.

### 2.7 HiC-LEGO captures stable early-stage and heterogeneous pleural-effusion chromatin organization in breast cancer

We analyzed the Hi-C compendium from van den Brand et al. [35], which includes healthy breast (HB), primary breast cancer (PB), liver metastasis (LM), and malignant pleural effusion (PE) samples. The study showed stable TAD and compartment organization across HB, PB, and LM but greater inter-patient heterogeneity in PE. We therefore focused on chromosome 13, where the study identified TAD heterogeneity among PE samples P09, P10, and P12 within chr13:50–75 Mb and stronger compartmentalization in P07 than P09 [35].

HiC-LEGO retained chromosome 13 contact organization beyond genomic-distance effects. Contact-like maps derived from reconstructed 3D distances correlated more strongly with the input Hi-C maps than genomic-distance-matched nulls in most samples (Fig. 5a; Supplementary Methods); we treated samples that did not exceed the null as lower-confidence reconstructions. Reconstructed TADs showed greater compactness, stronger 3D boundary separation, and smaller radii of gyration than size-matched controls (Fig. 5b), demonstrating that HiC-LEGO preserves domain-level spatial organization. HiC-LEGO partially recovered the reported P07–P09 compartment difference: four of five direct 3D metrics followed the expected direction, with stronger compartment segregation and neighborhood organization in P07 (Fig. 5c).

**Fig 5.**
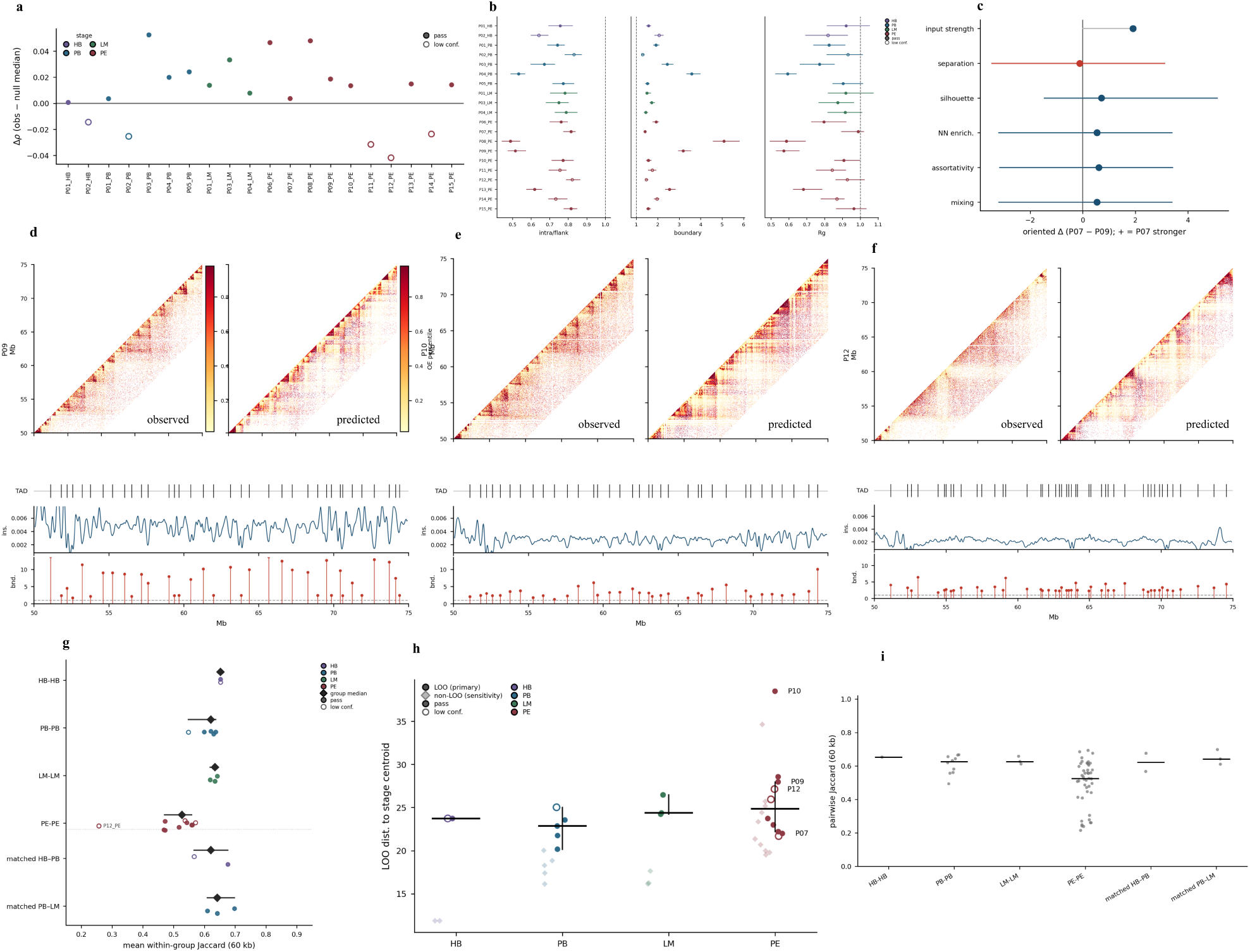
HiC-LEGO preserves disease-associated 3D genome organization across breast cancer progression. HB, healthy breast; PB, primary breast cancer; LM, liver metastasis; PE, pleural effusion. Filled points indicate samples exceeding the genomic-distance-matched null; open points indicate lowconfidence samples. (a) Most reconstructions show positive Δ*ρ*, indicating greater input–reconstruction concordance than expected under the matched null. (b) Input-defined TADs retain characteristic 3D organization relative to size-matched controls, with lower intra/flank distance and radius-of-gyration ratios and higher boundary scores (median *±* bootstrap 95% CI; dashed lines mark the null ratio of 1). (c) P07 shows stronger input compartmentalization than P09, with several direct-3D compartment metrics showing concordant trends; intervals denote null-derived ranges where available. (d–f) PE samples P09, P10, and P12 across chr13:50–75 Mb, showing correspondence between observed Hi-C and reconstructed contact-like maps and preservation of patient-specific TAD, insulation, and 3D boundary patterns (lag-wise log_2_(O/E), percentile-scaled; dashed boundary-score threshold = 1). (g) TAD-boundary similarity at 60 kb is lower and more heterogeneous within PE than within HB, PB, or LM, while patient-matched HB–PB and PB–LM pairs retain substantial boundary concordance (diamonds, group medians *±* bootstrap 95% CI). (h) Leaveone-out distance to stage centroids reveals greater structural heterogeneity among PE samples, including separation of P10; faint diamonds show non-LOO distances and black bars stage medians *±* bootstrap 95% CI. (i) Pairwise TAD-boundary Jaccard values show increased heterogeneity within PE and preserved boundary similarity in patient-matched cross-stage comparisons (black bars, group medians).

HiC-LEGO preserved patient-specific TAD organization across chr13:50–75 Mb (Fig. 5d–f). P09 showed clearer domain-associated contacts and 3D boundary signals, whereas P10 and P12 showed altered organization, consistent with the PE heterogeneity reported by van den Brand et al. [35]. The reconstructions preserved boundary positions and overall domain architecture more consistently than the reported P09*>*P10*>*P12 TAD-strength ordering. At a 60-kb boundary-matching tolerance, HB, PB, and LM retained substantial TAD-boundary overlap, and patient-matched HB–PB and PB–LM comparisons remained similar (Fig. 5g,i), supporting limited early-stage TAD reorganization. In contrast, PE samples showed greater dispersion in pairwise similarity, consistent with increased structural heterogeneity at this stage [35].

A leave-one-out analysis confirmed this pattern by comparing each sample with a stage-specific centroid derived from the remaining samples using boundary, insulation, and compartment profiles. PE reconstructions showed broader centroid-distance dispersion, with P07, P09, P10, and P12 occupying distinct positions relative to the PE centroid (Fig. 5h). Sensitivity analyses across boundary-matching tolerances, exclusion of lower-confidence reconstructions, and patient-matched versus unmatched comparisons produced similar overall trends (Supplementary Fig. S1). These results show that HiC-LEGO captures key breast cancer-associated 3D-genome features, including relative structural stability through HB, PB, and LM, patient-specific TAD and compartment organization, and increased architectural heterogeneity in malignant pleural effusions [35].

### 2.8 HiC-LEGO preserves breast cancer-associated TAD and compartment differences at 5-kb resolution

At 5-kb resolution, HiC-LEGO retained breast cancer-associated structural patterns reported by van den Brand et al. [35]. Fifteen of 20 chromosome 13 reconstructions exceeded the genomic-distance-matched null across HB, PB, LM, and PE samples (Fig. 6a); we treated the remaining five, including three PE samples, as lower confidence. Input-defined TADs showed smaller intra/flank distance ratios and radii of gyration and stronger 3D boundary separation than size-matched controls (Fig. 6b), showing that HiC-LEGO preserves domain-scale organization at 5-kb resolution. HiC-LEGO recovered the reported P07–P09 compartment difference: all five direct 3D metrics followed the expected direction and indicated stronger compartmentalization in P07 (Fig. 6c) [35].

**Fig 6.**
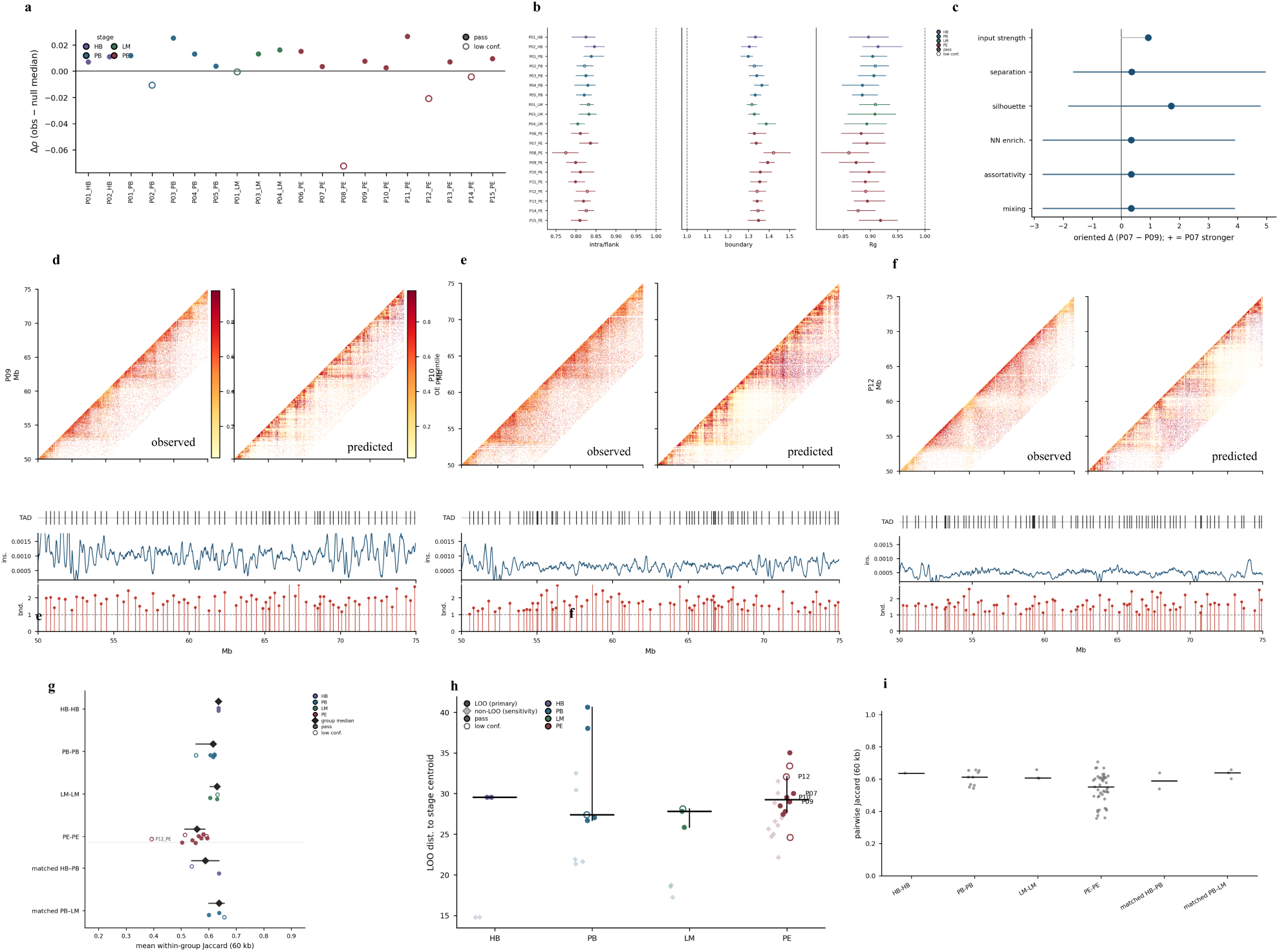
HiC-LEGO preserves disease-associated 3D genome organization at 5-kb resolution across breast cancer progression. HB, healthy breast; PB, primary breast cancer; LM, liver metastasis; PE, pleural effusion. Filled points indicate samples exceeding the genomic-distance-matched null; open points indicate low-confidence samples. (a) Most reconstructions show positive Δ*ρ*, indicating greater input–reconstruction concordance than expected under the matched null. (b) Input-defined TADs retain characteristic 3D organization relative to size-matched controls, with lower intra/flank distance and radiusof-gyration ratios and higher 3D boundary scores (median *±* bootstrap 95% CI; dashed lines denote a ratio of 1). (c) P07 shows stronger input compartmentalization than P09, with the direct-3D compartment metrics showing varying degrees of concordance; bars denote null-derived intervals where available. (d–f) PE samples P09, P10, and P12 across chr13:50–75 Mb show correspondence between observed Hi-C and reconstructed contact-like maps while retaining patient-specific TAD, insulation, and 3D boundary patterns (lag-wise log_2_(O/E), percentile-scaled; heatmaps block-averaged for display and tracks shown at native 5 kb; dashed boundary-score threshold = 1). (g) TAD-boundary similarity at 60 kb is lower and more heterogeneous within PE than within HB, PB, or LM, while patient-matched HB–PB and PB–LM comparisons retain substantial boundary concordance (diamonds, group medians *±* bootstrap 95% CI). (h) Leave-one-out distance to stage centroids, integrating master-boundary presence with boundary, intra/flank, insulation, and compartment profiles, reveals increased structural heterogeneity among PE samples; faint diamonds show non-LOO distances and black bars stage medians *±* bootstrap 95% CI. (i) Pairwise TAD-boundary Jaccard values show increased heterogeneity within PE and preservation of boundary similarity across patient-matched stage transitions (black bars,13group medians).

Within chr13:50–75 Mb, HiC-LEGO retained distinct patient-specific TAD organization across PE samples P09, P10, and P12 (Fig. 6d–f). P09 and P10 showed clearer domain organization, whereas P12 showed weaker, more fragmented patterns, consistent with previously reported PE heterogeneity [35]. Thus, HiCLEGO preserved sample-specific architecture without necessarily reproducing every reported TAD-strength difference. TAD-boundary comparisons showed relatively high similarity across HB, PB, and LM but lower and more variable similarity among PE samples (Fig. 6g,i). The stage-centroid leave-one-out analysis described above, which measures each sample’s multifeature distance from a centroid constructed from the remaining samples in the same stage, likewise showed greater PE variability (Fig. 6h). Resolution-specific sensitivity analyses and boundary-similarity comparisons produced the same overall trends (Supplementary Fig. S2). These results show that HiC-LEGO preserves key stage and patient-associated 3D-genome features at 5-kb resolution, including early-stage structural stability and increased heterogeneity in malignant pleural effusions [35].

### 2.9 HiC-LEGO captures multi-way chromatin proximity measured by Pore-C

We used experimentally observed GM12878 Pore-C multi-way contacts to independently validate higherorder chromatin organization in the 5-kb HiC-LEGO reconstructions [36, 37]. Native Pore-C contacts showed a clear span-dependent compaction relative to genomic-distance-matched controls (Fig. 7a). The effect emerged near the megabase scale and peaked at 1–5 Mb, where maximum pairwise 3D distance decreased by up to 10.2%, with similar trends for mean pairwise distance and radius of gyration. Short-range contacts below 100 kb showed little or no compaction, indicating that Pore-C provides the strongest higher-order validation at megabase scales.

**Fig 7.**
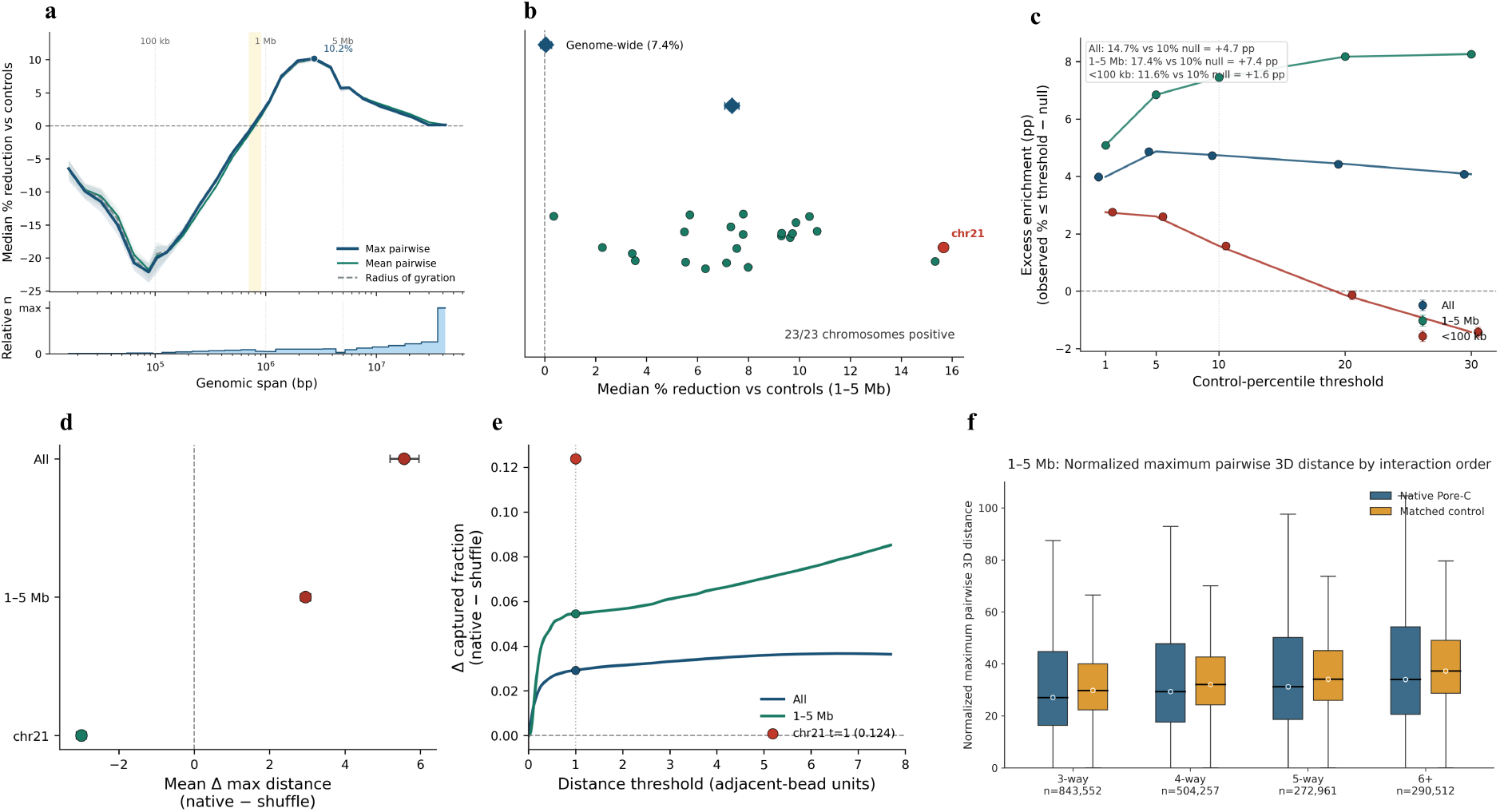
HiC-LEGO preserves Pore-C multi-way spatial compaction in GM12878. Native Pore-C concatemers contain *≥*3 *cis* anchors; genomic controls match chromosome, interaction order, and genomic gaps, and shuffle controls preserve native pairwise/genomic properties. 3D distances are normalized by chromosome-specific median adjacent-bead spacing. (a) Native interactions become more compact than matched controls at megabase scales, with maximum-pairwise distance reaching a 10.2% median reduction in the 1–5 Mb range (shading, bootstrap 95% CI). (b) This 1–5 Mb compaction is positive across all 23 chromosomes, with a genome-wide median reduction of 7.4%. (c) Native interactions are enriched in the compact tail of matched controls, particularly at 1–5 Mb (17.4% within the 10th control percentile versus 10% expected; +7.4 percentage points). (d,e) Comparison with anchor-shuffled controls shows scale- and chromosome-dependent differences in mean maximum distance, while native concatemers retain excess capture within compact-distance thresholds; chr21 shows a 0.124 increase in captured fraction at *t* = 1. (f) Native 1–5 Mb Pore-C interactions have lower normalized maximum pairwise 3D distances than matched genomic controls across 3-, 4-, 5-, and *≥*6-way interaction orders, supporting preservation of higher-order chromatin compaction.

Within 1–5 Mb, native contacts were more compact than matched controls across all 23 chromosomes, with a genome-wide interaction-weighted median reduction of 7.4% (Fig. 7b; Supplementary Fig. S3). Native contacts preferentially occupied the compact tail of the control distribution: 17.4% of 1–5 Mb contacts fell within the most compact 10% of controls, compared with the 10% null expectation (Fig. 7c). Under the stricter anchor-shuffle null [37], mean maximum distance did not favor native contacts genome-wide or across 1–5 Mb, although chromosome 21 showed clear compaction (Fig. 7d). Nevertheless, native concatemers occurred more frequently within compact 3D-distance thresholds than shuffled contacts, particularly at 1–5 Mb (Fig. 7e), supporting enrichment of compact higher-order configurations.

This signal extended across interaction complexity: native 3-, 4-, 5-, and *≥*6-way Pore-C contacts all showed lower normalized maximum pairwise distances than matched genomic controls within 1–5 Mb (Fig. 7f; *n* = 843552, 504257, 272961, and 290512, respectively). These results show that HiC-LEGO preserves experimentally observed multi-way chromatin organization, with the strongest and most consistent agreement at megabase-scale genomic spans.

## 3 Discussion

In this study, we developed HiC-LEGO, a domain-aware hierarchical framework for reconstructing highresolution 3D chromosome structures from chromatin contact maps. Rather than partitioning chromosomes solely by fixed genomic windows or relying on domains from a single caller, HiC-LEGO combines ensemble-based domain selection with graph-based local reconstruction and hierarchical assembly of domains into chromosome-scale structures (Fig. 1). Across five human cell lines at 5-kb resolution, HiC-LEGO achieved higher chromosome-scale reconstruction concordance than SOTA methods while better preserving TAD boundaries, boundary strength, and intra-domain organization (Fig. 2). These results indicate that the hierarchical strategy improves global reconstruction without sacrificing the local chromatin organization used to construct the model.

An important advantage of high-resolution 3D reconstruction is that the resulting coordinates provide a direct framework for examining spatial relationships that are difficult to interpret from contact maps alone. HiC-LEGO recovered experimentally supported FISH-probe relationships and compact CTCF/cohesin-associated loop anchors (Fig. 1f–k). At 1-kb resolution, the reconstructed GM12878 chromosome 8 placed an independently annotated *MYC* enhancer and promoter in close spatial proximity relative to genomic-distance-matched controls (Fig. 3). Similarly, Micro-C-derived structures retained finer local contact organization and spatial enrichment of experimentally defined microcompartment interactions (Fig. 4). These observations show that the hierarchical assembly preserves biologically relevant spatial information across scales ranging from regulatory interactions to complete chromosomes.

The reconstructed structures provide a means to examine chromatin organization that extends beyond individual pairwise contacts. Across breast cancer progression, HiC-LEGO retained broad differences in TAD, compartment, and inter-patient organization reported in the experimental data, including increased structural heterogeneity among malignant pleural effusion samples (Figs. 5 and 6). These patterns were recovered more consistently at the level of overall domain and compartment organization than as exact quantitative differences in every individual feature. Pore-C provided a complementary higher-order test: experimentally observed multi-way contacts were enriched in compact reconstructed configurations relative to matched genomic controls, with the strongest signal at megabase-scale genomic spans (Fig. 7). The stricter pairwise/genomic-preserving shuffle did not show a uniform native compactness advantage in mean distance, but retained enrichment within compact-distance thresholds. Thus, the reconstructed coordinates capture higher-order spatial organization beyond simple chromosome-wide contact concordance, while remaining appropriately interpreted as structures constrained primarily by pairwise contact information.

Overall, HiC-LEGO provides a modular framework for converting high-resolution chromatin contact data into interpretable chromosome-scale 3D models while retaining biologically defined domain organization. Its agreement with contact-map structure, regulatory interactions, Micro-C features, disease-associated chromatin organization, and multi-way Pore-C contacts supports the use of these reconstructions as structural references for integrating genomic and epigenomic information.

## 4 Methods

### 4.1 Ensemble-based TAD detection for robust domain identification

TADs provide biologically motivated structural units for hierarchical 3D genome reconstruction [2]. However, TAD-calling methods use different definitions and statistical assumptions and can therefore produce substantially different domain partitions from the same Hi-C map [38–40]. Reliance on a single caller may consequently propagate method-specific boundary errors into downstream reconstruction. HiC-LEGO addresses this variability by integrating domain predictions from multiple TAD callers and selecting a structurally concordant domain set.

We quantifies the concordance between two domain sets, *A* and *B*, using the Measure of Concordance (MoC) [23]:

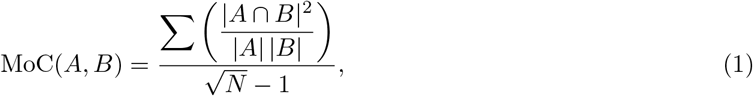

where |*A ∩ B*| is the length of the overlap between two domains, |*A*| and |*B*| are their respective lengths, and *N* denotes the number of comparisons. Thus, Eq. 1 assigns higher scores to domain sets with greater normalized interval overlap. We evaluated three strategies for deriving an optimal domain set from the ensemble (Fig. 8).

**Fig 8.**
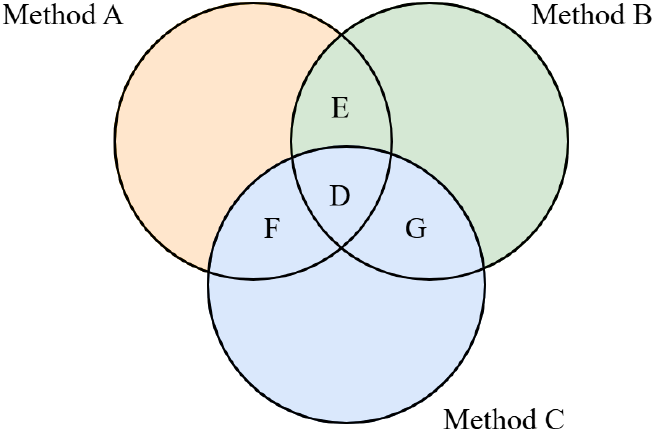
Strategies evaluated for selecting an optimal domain set from multiple TAD callers. *A, B*, and *C* denote domain sets produced by three independent TAD-calling methods.

#### Hypothesis 1: complete cross-method agreement

Only domains supported by all available TAD callers were retained. For three domain sets *A, B*, and *C*, this corresponds conceptually to the common overlap *A ∩ B ∩ C* (region *D* in Fig. 8). This stringent strategy retains only domains with non-zero overlap across every method.

#### Hypothesis 2: pairwise structural concordance

All pairwise MoC scores between TAD callers were first calculated using Eq. 1, and we selected the pair of domain sets with the highest concordance. For the selected pair, we compared each domain from the first set with all domains in the second set, and a domain is retained if it has a non-zero MoC with at least one domain in the second set. This procedure removes domains unsupported by the most concordant independent caller while avoiding the stringent requirement of agreement across all methods. In Fig. 8, the candidate pairwise overlaps correspond to *A ∩ B, A ∩ C*, and *B ∩ C*.

#### Hypothesis 3: one-versus-all concordance

For each domain set *X* in the ensemble, we calculated its average MoC against all other domain sets:

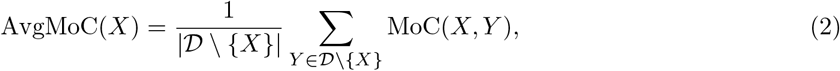

where *D* denotes the complete collection of TAD-call sets. Optimal domain set is selected as

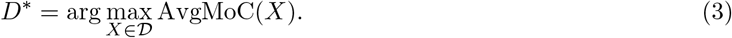

Thus, Eqs. 2 and 3 select the single TAD-call set with the greatest average structural agreement with the remaining methods. We evealuated the three strategies by their effect on downstream 3D reconstruction, and used the best-performing strategy the final HiC-LEGO pipeline.

### 4.2 HiC-LEGO reconstruction framework and input processing

HiC-LEGO reconstructs high-resolution chromosome structures through a hierarchical process that combines domain-level graph modeling, fine-resolution reconstruction of individual domains, megabase-scale assembly, and chromosome-wide refinement. For each chromosome, the reconstruction uses a fine-resolution Hi-C interaction matrix, the selected chromatin domains, and a corresponding 1-Mb Hi-C interaction matrix. Hi-C matrices at the required resolutions were extracted and Knight-Ruiz (KR) [41] normalized using Juicer Tools [42]. Chromatin domains were represented as half-open genomic intervals [*s*_*k*_, *e*_*k*_).

The reconstruction proceeds at three structural levels. First, fine-resolution Hi-C contacts are aggregated between selected domains to construct a chromosome-specific domain graph. A graph neural network (GNN) is trained on this graph to infer the spatial organization of the domains, and the learned representation and network parameters are transferred to the fine-resolution genomic bins within each domain. Second, the independently reconstructed domains are assembled into megabase-indexed LEGO regions using fine-resolution Hi-C constraints. Third, a chromosome-scale backbone is independently reconstructed from the corresponding 1-Mb Hi-C matrix using the same graph-based structural inference architecture. The assembled LEGO regions are positioned along this backbone and refined using selected fine-resolution intra- and inter-region Hi-C contacts.

### 4.3 Graph-based reconstruction of chromatin domains

For two retained domains *D*_*a*_ and *D*_*b*_, with corresponding sets of fine-resolution genomic bins ℬ_*a*_ and ℬ_*b*_, respectively, the domain–domain interaction is calculated as

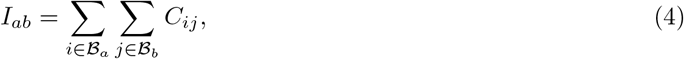

where *C*_*ij*_ denotes the KR-normalized Hi-C interaction frequency between genomic bins *i* and *j*, and *I*_*ab*_ is the total interaction between domains *a* and *b*, hence, Eq. 4 representing an interaction sum. Domains without mapped genomic coordinates or non-zero aggregate interactions are excluded before reconstruction. Each retained domain is represented by one graph node. The domain–domain interaction matrix is KR-balanced for graph construction. Each node is initialized with a 512-dimensional second-order LINE embedding [43] learned from the aggregated domain interaction matrix. LINE embeddings are trained for 100 epochs using Adam optimizer [44] with a learning rate of 0.01, batch size 4096, and a negative-sampling ratio of 5.

The structural inference model uses mean-neighbour aggregation together with a separate transformation of the representation of the node itself,

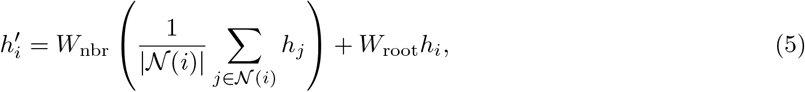

where *h*_*i*_ is the representation of node *i, N*(*i*) is its set of neighbouring nodes, and *W*_nbr_ and *W*_root_ are learned linear transformations. Equation 5 combines the mean representation of neighbouring domains with a separately transformed representation of the domain itself. The graph layer maps 512 input features to 512 features and is followed by fully connected layers with dimensions 512 *→* 256 *→* 128 *→* 64 *→* 3. ReLU activation is applied after each hidden layer, whereas the final three-dimensional output has no activation and represents the Cartesian coordinate *x*_*i*_ *∈* ℝ^3^ of each domain.

Since higher Hi-C interaction frequencies are expected to correspond to smaller spatial separations, domain interactions are converted to target distances using an inverse power-law relationship,

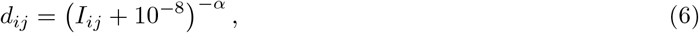

where *I*_*ij*_ is the aggregated domain–domain interaction before the additional KR balancing used for graph construction, *d*_*ij*_ is the corresponding target spatial distance, and *α >* 0 is conversion factor. The constant 10^−8^ in Eq. 6 is included only for numerical stability to prevent singular values for zero or nearzero interactions and has no physical interpretation. The GNN predicts pairwise Euclidean distances 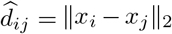 from the inferred coordinates. For each candidate value of *α*, the model is trained by minimizing

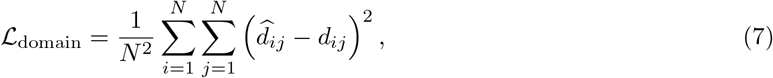

where *N* is the number of retained domains. Equation 7 aligns the Euclidean distances in the predicted structure towards the distances implied by the Hi-C interaction frequencies.

Since the appropriate contact-to-distance relationship is not known while starting, a grid search s *α ∈ {*0.1, 0.2, …, 1.9} is implemented. For each, the network is optimized using Adam with a learning rate of 10^−3^ at most 1000 epochs, with convergence defined by an absolute change in loss below 10^−8^. The value of *α* producing the highest Spearman correlation between the reconstructed and target-distance matrices is chosen as the optimal one. A separate model is trained for each chromosome. The chromosome-level representation is transferred to the fine-resolution bins within each domain. For domain *D*_*k*_ containing *n*_*k*_ genomic bins, the corresponding fine-resolution Hi-C submatrix is extracted and KR-balanced, and a new 512-dimensional LINE embedding 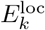 is learned from the local interaction matrix. The chromosome-level domain embedding *e*_*k*_ is replicated across the bins in the domain, and the local embedding is aligned to this representation using orthogonal Procrustes transformation [45],

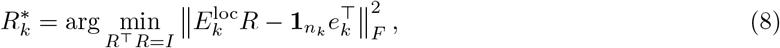

where 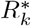 is the optimal orthogonal transformation and 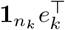 denotes the chromosome-level domain embedding replicated across the *n*_*k*_ fine-resolution bins. The aligned embedding 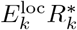 from Eq. 8 is supplied to the chromosome-trained network. The chromosome-level model parameters are loaded as initialization and fine-tuned independently for each domain rather than being held fixed.

Within each domain, KR-normalized interaction frequencies are bounded below by 10^−12^ for numerical stability and converted to target distances using the same inverse-power relationship. Candidate exponents *α ∈ {*0.1, 0.2, …, 2.0} are evaluated separately for each domain. Fine-tuning minimizes the upper-triangular pairwise distance error,

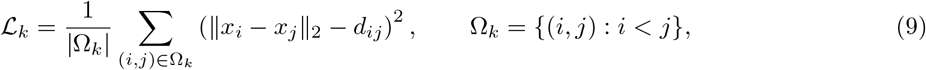

where Ω_*k*_ denotes the set of unique bin pairs within domain *k*. Equation 9 is optimized until the absolute change in loss falls below 10^−4^, and the exponent producing the highest Spearman distance correlation is retained as the optimal conversion factor for that domain.

### 4.4 Hierarchical assembly of domains and chromosome-scale reconstruction

The independently reconstructed domains are then grouped into megabase-indexed LEGO regions according to the 1-Mb genomic interval containing the start coordinate of each domain. Domains extending across a 1-Mb boundary are retained intact in the region where they started rather than being split. A fine-resolution genomic coordinate lattice is constructed for each LEGO region, and the reconstructed domain coordinates are inserted at their corresponding genomic positions.

Within each LEGO region, positive fine-resolution Hi-C interactions are converted to target spatial distances using 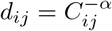, whereas zero-valued interactions are excluded. The conversion exponent for each LEGO region is set to the median of the optimal *α* values obtained from the domains it constitutes of.

Each reconstructed domain structure *X*_*k*_ is allowed an isotropic scale, three-dimensional rotation, and translation,

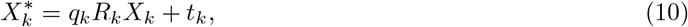

where *q*_*k*_ is the isotropic scale factor, *R*_*k*_ is the rotation matrix, and *t*_*k*_ *∈* ℝ^3^ is the translation vector. The transformation in Eq. 10 allows independently reconstructed domains to be assembled while preserving their internal relative geometry apart from a uniform scale.

Domains are optimized sequentially by coordinate descent to maximize agreement between pairwise distances in the assembled structure and those implied by the fine-resolution Hi-C data. For the set *P* of unique bin pairs with positive Hi-C interactions within a LEGO region, the optimization criterion is

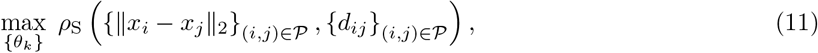

where *ρ*_S_ denotes Spearman correlation and *θ*_*k*_ represents the scale, rotation, and translation parameters of domain *k*. Thus, Eq. 11 maximizes rank agreement between distances in the assembled structure and the corresponding Hi-C-derived target distances. The reconstruction with the highest distance correlation is kept. Fine-resolution genomic bins lying between reconstructed domains are incorporated after domain assembly while the coordinates of the reconstructed domains remain fixed. Gap coordinates are optimized by maximizing the bounded Lorentzian similarity

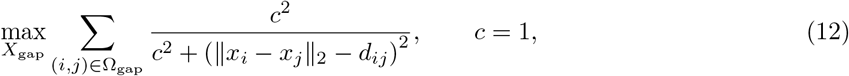

where Ω_gap_ contains informative Hi-C pairs including gap coordinates. Equation 12 helps reconstructed distances that agree with the Hi-C-derived targets while bounding the influence of large residuals. A bead-level refinement regularizes local smoothness and genomic-chain continuity.

For chromosome-scale organization, HiC-LEGO independently reconstructs a low-resolution chromosome backbone from the corresponding 1-Mb KR-normalized Hi-C matrix using the same GNN architecture and contact-to-distance fitting strategy described above. The resulting 1-Mb coordinates act as a backbone for assembly of the fine-resolution LEGO regions. Each LEGO region is associated with the backbone position corresponding to its midpoint and is translated so that its centroid coincides with the corresponding backbone coordinate.

Chromosome-wide refinement uses a selected subset of fine-resolution Hi-C contacts rather than all possible chromosome-wide pairs. Inter-region contacts are selected primarily between neighbouring and next-neighbour LEGO regions using genomic-distance stratification and interaction-frequency ranking, whereas the number of intra-region contacts is capped to prevent densely connected local regions from dominating the global optimization.

For each retained contact (*u, v*), an unscaled contact-derived distance is defined as

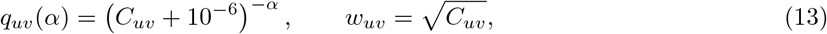

where *C*_*u*V_ is the fine-resolution KR-normalized Hi-C interaction frequency, *α* is the conversion factor, and 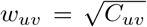 provides sublinear weighting that gives stronger Hi-C contacts greater influence while limiting dominance by the highest interaction frequencies. The 10^−6^ term in Eq. 13 is included for numerical stability.

For each candidate *α ∈* {0.1, 0.2, …, 2.0}, scaling factor is estimated as:

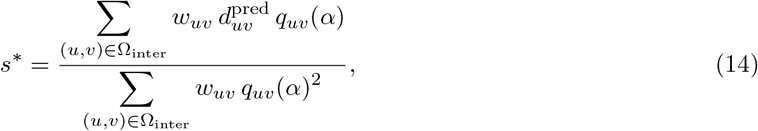

where 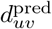 is the current Euclidean distance between bins *u* and *v*, and Ω_inter_ denotes the selected inter-region contact set. Equation 14 provides a closed-form weighted least-squares estimate of the global distance scale. The candidate exponent yielding the lowest Huber-weighted inter-region residual is retained, and the 1-Mb backbone is rescaled by the corresponding *s*\*. The chromosome-scale target distance for each selected pair is therefore 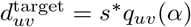.

After initial backbone placement, each LEGO region is refined as a rigid body using an axis–angle rotation and translation. For LEGO region *k*, the optimization objective is

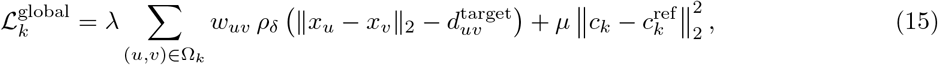

where Ω_*k*_ contains the selected Hi-C contacts involving region *k, ρ*_*δ*_ is the Huber loss [46] with threshold *δ, c*_*k*_ is the current centroid of the LEGO region, and 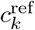 is its backbone-derived reference centroid. The first term in Eq. 15 promotes agreement with fine-resolution Hi-C-derived spatial constraints, whereas the second penalizes large departures from the chromosome-scale backbone.

LEGO regions are updated step-by-step in Gauss–Seidel order using L-BFGS-B [47]. After the first rigid refinement, the fine-resolution bins not present in an assembled LEGO structure are put into the chromosome space using spline interpolation together with boundary alignment. The system then recalculates the contact-to-distance relationship to perform a second coarse alignment using inter and intra contacts. Throughout this refinement, the LEGO blocks are optimized by adjusting their overall position and orientation as rigid units.

### 4.5 Computational implementation and structural outputs

HiC-LEGO is implemented in Python using PyTorch, SciPy, and Numba. LINE embedding, GNN training, and fine-resolution domain reconstruction use GPU when available. Graph models are trained independently for each chromosome, and the chromosome-level model state is transferred only to the corresponding fine-resolution domains. The final chromosome reconstruction contains one Cartesian coordinate for each valid fine-resolution genomic bin represented in the coordinate mapping. Structural coordinates are expressed in arbitrary model units and are not calibrated to physical nanometres or angstroms. PDB files are generated for structural visualization.

## Data availability

Hi-C contact maps used to reconstruct chromosome structures for GM12878, HUVEC, K562, HMEC, and IMR90 are available under GEO accession GSE63525. HiCCUPS chromatin loops with CTCF motif annotations and GM12878 subcompartment annotations from the same series were used for loop and compartment validation.

Breast cancer progression Hi-C files are available under GEO accession GSE273998. Genome-wide mouse embryonic stem cell (mESC) Micro-C data are available under GSE130275, Region Capture Micro-C (RCMC) data under GSE207225, and Bonev et al. mESC Hi-C data under GSE96107.

GM12878 Pore-C (NlaIII) multi-way fragment alignments and pairwise cooler matrices are available under GEO accession GSE149117, including sample GSM4490689.

ENCODE ChIP-seq and chromatin-state datasets used in this study include GM12878 CTCF (wgEncodeAwgTfbsBroadGm12878CtcfUniPk), RAD21 (wgEncodeAwgTfbsHaibGm12878Rad21V0416101UniPk), H3K27ac (GSM733771), and ChromHMM annotations (GSM936082). mESC annotation datasets are available under GEO for CTCF (GSE90994), and through ENCODE for H3K4me1 peaks (ENCFF763GNF). Gene annotations were obtained from GENCODE human releases v19 and v38, and mouse release vM38.

The final output pdb files including 1kb structure and 5kb structures from all the experiments are in our zenodo repository.

## Code availability

The HiC-LEGO tool is available at https://github.com/OluwadareLab/HiC-LEGO.

## Acknowledgments

We would also like to thank all the members of OluwadareLab for their inputs and comments during discussions.

