## Supplemental File 1 for "HiC-LEGO: Biologically Guided High-Resolution 3D Genome Reconstruction Preserves Chromatin Organization at Kilobase Resolution"

#### 1 Supplementary Methods

##### 1.1 CTCF/RAD21 loop-anchor validation

We used intra-chromosomal GM12878 HiCCUPS loops from Rao et al. [1] for CTCF validation. Starting from 9,448 loops, loops with either anchor overlapping blacklist or gap regions were removed, leaving 9,321 loops. We then retained loops with CTCF signal at both anchors (4,990), followed by RAD21 signal at both anchors (4,645). Of these, 4,644 mapped to distinct beads in the 5-kb HiC-LEGO reconstructions. The analysis shown in Fig. 1j was restricted to autosomes, resulting in 4,514 CTCF+RAD21-supported loops (Supplementary Table S2).

We calculated the Euclidean distance between the reconstructed 3D coordinates of the two loop-anchor beads for each retained loop. We compared each loop with 100 genomic-distance-matched CTCF-random control pairs. Across the autosomal loop set, the median reconstructed anchor distance was approximately 38.6 model units, compared with approximately 47.6 for the matched controls (Wilcoxon rank-sum $p = 7.46 \times 10^{-45}$). No per-loop significance or percentile threshold was used to select loops for this analysis. CTCF motif orientation was not used as a retention criterion.

The representative loop shown in Fig. 1k connects chr22:23,880,000–23,890,000 and chr22:24,100,000–24,110,000, spanning 220 kb. Its reconstructed anchor distance was 1.517 model units, compared with a median matched-control distance of 57.76 model units (Supplementary Table S2).

##### 1.2 Micro-C and microcompartment validation

We obtained Wild-type mouse embryonic stem-cell (mESC) Micro-C data from GEO accession GSE130275 [2]. We extracted Micro-C contacts at 5-kb resolution using Juicer `dump observed KR` and we obtained the corresponding mESC Hi-C comparator was obtained from Bonev et al. [3] (GEO accession GSE96107). Four embryonic stem-cell Hi-C replicates were binned at 5 kb, pooled, and KR/ICE-balanced before reconstruction. HiC-LEGO structures were reconstructed independently from the Micro-C and Hi-C contact matrices.

Chromosomes 3, 5, 8, and 18 were analyzed because they contain the RCMC capture regions for Sox2, Ppm1g, Klf1, and Fbn2, respectively, for which corresponding chromosome-scale Micro-C reconstructions were available. The genome-wide Micro-C dataset itself was not restricted to these chromosomes.

For microcompartment validation, we used the published RCMC BEDPE interaction set from Goel et al. [4]. The Ppm1g analysis was restricted to the 249 cis interaction pairs on chromosome 5 within the RCMC capture region chr5:31,257,344–32,382,344 (mm39). For each BEDPE anchor, the interval midpoint was mapped to the containing 5-kb reconstructed bin; when no containing bin was available, the nearest reconstructed bin center was used. Interactions involving missing reconstructed coordinates were omitted. The Euclidean distance between the two mapped anchor coordinates was calculated as

$$d_{ij} = \sqrt{(x_i - x_j)^2 + (y_i - y_j)^2 + (z_i - z_j)^2}. \tag{S1}$$

For each of the 249 microcompartment interactions, 50 background bin pairs were sampled from non-loop pairs within the Ppm1g region. Background pairs were matched to the genomic separation of the corresponding interaction using a tolerance of $\pm \max(10 \text{ kb}, 0.15 d_{\text{genomic}})$. Known loop bin pairs were excluded from the candidate background pool. Random sampling was reproducibly seeded independently for each interaction. This procedure yielded 12,450 matched background pairs.

The median reconstructed 3D distance was 12.751 for the annotated microcompartment interactions and 15.785 for the matched background pairs, corresponding to a 1.238-fold enrichment in spatial proximity. The reported significance was calculated using a one-sided Mann–Whitney U test comparing the 249 interaction distances with the pooled 12,450 background distances ($U = 1{,}199{,}841$, $p = 4.87 \times 10^{-10}$).

For 3D aggregate peak analysis, reconstructed pairwise distances were converted to a contact-like matrix according to

$$C_{ij}^{3D} = \frac{1}{d_{ij} + 1}. \tag{S2}$$

The diagonal was excluded. For each of the 249 mapped interactions, a ±5-bin (±25-kb) window was extracted and the resulting patches were averaged. The 3D aggregate peak analysis (APA) score was defined as the center value divided by the mean of all pixels outside the central $3 \times 3$ region. This analysis produced a 3D APA score of 1.035, reported as 1.04 in Fig. 4.

#### 1.3 Breast cancer progression validation

**Dataset and reconstruction inputs.** Breast cancer progression analyses used the Hi-C compendium of van den Brand et al. [5]. Breast cancer progression analyses were performed on chromosome 13 at 20-kb resolution using 20 samples spanning healthy breast (HB), primary breast cancer (PB), liver metastasis (LM), and malignant pleural effusion (PE) stages. All analyses used the hg38 genome assembly. The supplied HiC-LEGO domain sets were reused throughout the validation experiments. Sample identities and stage assignments are provided in Supplementary Table S3.

**Chromosome-scale reconstruction concordance and genomic-distance null.** For each reconstructed chromosome, pairwise Euclidean distances were converted to contact values according to

$$C_{ij}^{3D} = \frac{1}{(d_{ij} + \epsilon)^{\alpha}}, \qquad \epsilon = 10^{-6}, \tag{S3}$$

where $\alpha$ is conversion factor from the HiC-LEGO reconstruction. Diagonal entries were excluded. Spearman correlation was calculated between the input Hi-C matrix and reconstructed contact-like matrix using upper-triangular pairs with finite positive Hi-C contacts. A genomic-distance-matched null was generated by shuffling reconstructed contact values within genomic-distance strata while preserving the genomic-distance distribution. Reconstruction improvement over the null was quantified as

$$\Delta\rho = \rho_{\text{real}} - \text{median}\left(\rho_{\text{null}}\right). \tag{S4}$$

Samples were classified as passing the null comparison only when the observed correlation exceeded both the median and the 95th percentile of the genomic-distance-matched null distribution. Samples failing this criterion were displayed as lower-confidence reconstructions.

**TAD structural validation.** Structural preservation of the supplied TADs was evaluated using three direct 3D measures. TAD compactness was quantified as the mean pairwise 3D distance within the TAD divided by the mean distance to equal-bin-count flanking regions. The 3D boundary score was calculated over a fixed 100-kb window (five 20-kb bins) as

$$B_{3D} = \frac{\text{mean}\, d_{\text{cross}}(L, R)}{\text{mean}\left[d_{\text{within}}(L), d_{\text{within}}(R)\right]}, \tag{S5}$$

where $L$ and $R$ denote bins on opposite sides of a TAD boundary. Scores from the left and right edges of each TAD were averaged. Radius of gyration was calculated as

$$R_g = \sqrt{\frac{1}{N}\sum_{i=1}^{N} \|\mathbf{x}_i - \bar{\mathbf{x}}\|_2^2}. \tag{S6}$$

Each TAD was compared with up to 20 genomic control regions having the same number of bins and a genomic span within 15% of the corresponding TAD. Controls overlapping excluded genomic regions, including gaps, blacklist regions, and centromeric regions, were omitted. For each metric, the real value was divided by the median of its matched controls, and the plotted sample-level value represents the median of these per-TAD ratios. Bootstrap 95% confidence intervals were obtained from 1,000 percentile-bootstrap resamples.

**Compartment validation.** A/B compartment assignments were derived from the input Hi-C matrix. Observed contacts were normalized by their mean at each genomic separation, the resulting observed/expected matrix was centered by subtracting one, values were clipped at the 99.5th percentile, and the leading eigenvector was calculated. Eigenvector sign was oriented such that its Spearman correlation with GC content was non-negative; bins with non-negative eigenvector values were assigned to compartment A and remaining bins to compartment B. Direct 3D compartment preservation was evaluated using five complementary measures: A/B spatial separation, silhouette score, nearest-neighbor enrichment, graph assortativity, and compartment mixing. Spatial separation was defined as the mean A–B distance divided by the mean of the within-A and within-B distances. Silhouette scores were calculated from Euclidean 3D coordinates. Nearest-neighbor enrichment was calculated as the observed fraction of same-compartment neighbors relative to its random expectation using $k = 15$ nearest neighbors. Assortativity was calculated on the corresponding nearest-neighbor graph, while mixing was defined from the fraction of neighboring beads assigned to different compartments. Null distributions were generated by block-preserving circular shifts of the A/B compartment assignments using 120 permutations with random seed 42. Each direct 3D metric was standardized relative to its null distribution as

$$Z_{\text{metric}} = \frac{M_{\text{obs}} - \text{median}(M_{\text{null}})}{\text{SD}(M_{\text{null}})}, \tag{S7}$$

with the sign of the mixing metric inverted so that larger values consistently indicate stronger compartmentalization. The P07–P09 comparison was calculated as the difference between these oriented effects. Input compartment strength was compared using cohort-standardized compartment scores. Approximate uncertainty intervals shown in the figure were derived from the corresponding null 5th and 95th percentiles.

**Regional TAD analysis.** Patient-specific domain organization was evaluated within chr13:50–75 Mb for PE samples P09, P10, and P12. Observed and reconstructed contact-like matrices were transformed using lag-wise $\log_2$(observed/expected) normalization for genomic separations up to 10 Mb. The input Hi-C matrix was lightly Gaussian-smoothed ($\sigma = 0.8$), and each matrix was independently percentile-scaled for visualization between the 2nd and 98th percentiles. Insulation was calculated from the input Hi-C matrix using the mean interaction across a 200-kb left–right window, excluding the nearest 40 kb around the diagonal, followed by Gaussian smoothing with $\sigma = 1.5$ bins. Supplied TAD boundaries overlapping the region were displayed directly, while reconstructed 3D boundary scores were calculated using the same fixed-window definition described above.

**TAD-boundary similarity across disease stages.** Boundary similarity was evaluated with a 60-kb matching tolerance. Boundaries from two samples were greedily matched one-to-one to their nearest counterpart when their genomic positions differed by no more than 60 kb. Boundary similarity was then calculated using the Jaccard index,

$$J(A, B) = \frac{|A \cap B|}{|A \cup B|}. \tag{S8}$$

Within-stage comparisons included HB–HB, PB–PB, LM–LM, and PE–PE sample pairs. Patient-matched cross-stage comparisons included P01 and P02 for HB–PB and P01, P03, and P04 for PB–LM. For the summarized analysis, each sample was represented by its mean Jaccard similarity to the other samples in the same comparison group. Group medians and 95% confidence intervals were estimated using 2,000 bootstrap resamples.

**Leave-one-out stage heterogeneity.** Stage-specific structural heterogeneity was quantified using a concatenated feature representation comprising master-boundary presence, 3D boundary scores, intra/flank ratios, insulation values at master boundaries, and compartment eigenvector profiles. Boundary-score, intra/flank, insulation, and compartment features were standardized within each sample; binary master-boundary presence was retained without z-scoring. For sample $i$ within a disease stage, the leave-one-out stage centroid was defined as the mean feature vector of all remaining samples in the same stage,

$$\bar{\mathbf{x}}_{-i} = \frac{1}{n-1} \sum_{j \neq i} \mathbf{x}_j, \tag{S9}$$

and structural deviation was measured as

$$d_i^{\text{LOO}} = \|\mathbf{x}_i - \bar{\mathbf{x}}_{-i}\|_2 \,. \tag{S10}$$

The non-leave-one-out values shown as secondary points were calculated relative to the centroid containing all samples from the corresponding stage. Stage-level medians and 95% confidence intervals were obtained from 2,000 bootstrap resamples of the leave-one-out distances.

#### 1.4 Breast cancer progression validation at 5-kb resolution

The 5-kb breast cancer analysis used the same 20-sample cohort, chromosome 13 validation framework, and hg38 genome assembly described for the 20-kb analysis (Section 1.3). Sample identities and disease-stage assignments were unchanged (Supplementary Table S3). Input Hi-C matrices and reconstructed structures were analyzed at 5-kb resolution. The reconstruction domain sets were the corresponding 5-kb supplied HiC-LEGO TAD lists.

Chromosome-scale reconstruction concordance was evaluated using the same procedure as for the 20-kb structures. Pairwise 3D distances were converted to contact values using Eq. S3, with the sample-specific final HiC-LEGO exponent retained without re-fitting. Spearman correlation was calculated over finite, positive upper-triangular Hi-C entries separated by at least one bin, with at most $2 \times 10^6$ pairs evaluated per sample. The genomic-distance-matched null consisted of 10 shuffles stratified by 5-kb genomic-separation bins. A reconstruction was classified as passing only when its observed correlation exceeded both the median and the 95th percentile of the corresponding null distribution.

Supplied-TAD preservation was evaluated using the same intra/flank distance ratio, direct 3D boundary score, and radius-of-gyration metrics defined for the 20-kb analysis. Eligible TADs contained at least three reconstructed bins, spanned at least 60 kb, and had reconstruction coverage of at least 0.5. Each TAD was compared with up to 20 genomic controls matched for bin count and genomic span within 15%, excluding gaps, blacklist regions, centromeric regions, and other excluded intervals. The direct 3D boundary score used the same fixed physical window of 100 kb, corresponding to 20 bins at 5-kb resolution. Sample-level real/control ratios and percentile-bootstrap 95% confidence intervals were calculated as described for the 20-kb analysis using 1,000 bootstrap resamples.

A/B compartment analysis used the same observed/expected normalization, leading-eigenvector assignment, GC-based eigenvector orientation, and five direct 3D compartment metrics described in Section 1.3. Nearest-neighbor enrichment, mixing, and assortativity were all evaluated on a $k = 15$ nearest-neighbor graph. Null distributions were generated using 120 block-preserving circular shifts of compartment labels. The P07–P09 comparison was calculated from the oriented null-standardized 3D effects, whereas input compartment strength was represented by the cohort-standardized input Hi-C compartment score.

Regional validation was performed within chr13:50–75 Mb for PE samples P09, P10, and P12. Hi-C insulation was calculated using a 200-kb window with a 40-kb diagonal exclusion and Gaussian smoothing of $\sigma = 1.5$ bins. Supplied TAD boundaries and direct 3D boundary scores were evaluated at the native 5-kb resolution. For visualization, the regional Hi-C and reconstructed contact-like heatmaps were block-averaged, whereas all quantitative tracks and structural measurements were calculated from the native 5-kb data.

TAD-boundary similarity and stage heterogeneity were evaluated using the same procedures as at 20 kb. Boundary Jaccard similarity used greedy one-to-one matching within a 60-kb tolerance, with 20- and 40-kb tolerances additionally examined as sensitivity analyses (Supplementary Fig. S2). Leave-one-out stage heterogeneity was calculated from a concatenated feature vector containing master-boundary presence together with within-sample standardized 3D boundary scores, intra/flank ratios, insulation values, and compartment eigenvector profiles. Euclidean distance to the remaining-sample stage centroid was used as the primary heterogeneity measure.

#### 1.5 Pore-C multi-way validation

GM12878 NlaIII Pore-C fragment alignments (GSM4490689) were processed across 13 pooled runs [6, 7]. GRCh38-aligned fragments were lifted to hg19 for mapping to the HiC-LEGO structures. Valid *cis* concatemers were required to map to a single chromosome, retain at least 60% of their original fragments, and contain at least three unique 5-kb reconstructed beads after collapsing multiple fragments mapping to the same bead. This yielded 11,639,205 validation-ready concatemers, of which 11,635,680 had at least 10 valid matched controls and were used for the primary analyses.

Matched genomic controls were generated on the same chromosome with the same interaction order and approximately preserved consecutive genomic-gap structure. Relative gap and genomic-span errors were required to be $\leq 0.35$. Twenty controls were targeted per native concatemer, and concatemers with fewer than 10 valid controls were excluded. For each compactness metric, the median of the accepted controls was used as the matched-control value. Compactness was quantified using maximum pairwise 3D distance, mean pairwise 3D distance, and radius of gyration. Distances were additionally normalized by the chromosome-specific median Euclidean distance between adjacent reconstructed beads.

Percent reduction relative to matched controls was calculated as

$$\text{Reduction} = 100\left(1 - \frac{d_{\text{native}}}{d_{\text{control}}}\right), \tag{S11}$$

where $d_{\text{control}}$ is the median value across matched controls. Genomic-span effects were evaluated in log-spaced bins, retaining bins with at least 200 concatemers. The 1–5 Mb analysis included concatemers with 1 Mb $\leq$ span $<$ 5 Mb. Median effects and 95% confidence intervals were estimated by nonparametric bootstrap.

For compact-tail enrichment, each native concatemer was assigned a percentile relative to its matched-control maximum pairwise distances. Enrichment was evaluated at the 1st, 5th, 10th, 20th, and 30th control percentiles and reported as the observed fraction minus the corresponding uniform-null expectation.

A stricter shuffle control was additionally generated by rearranging anchors on the same chromosome while preserving interaction order, genomic-gap and span characteristics, and approximately preserving pairwise contact properties, thereby disrupting the original multi-way anchor combination. Twenty shuffled realizations were targeted per native concatemer. Native maximum pairwise distances were compared with the median accepted shuffled distance after adjacent-bead normalization. Captured fraction at threshold $t$ was defined as the proportion of concatemers with normalized maximum pairwise distance $\leq t$.

Interaction-order analyses used the effective cardinality after 5-kb mapping and duplicate-bead collapse, with concatemers grouped as 3-way, 4-way, 5-way, or $\geq$ 6-way.

### 2 Supplementary Figures

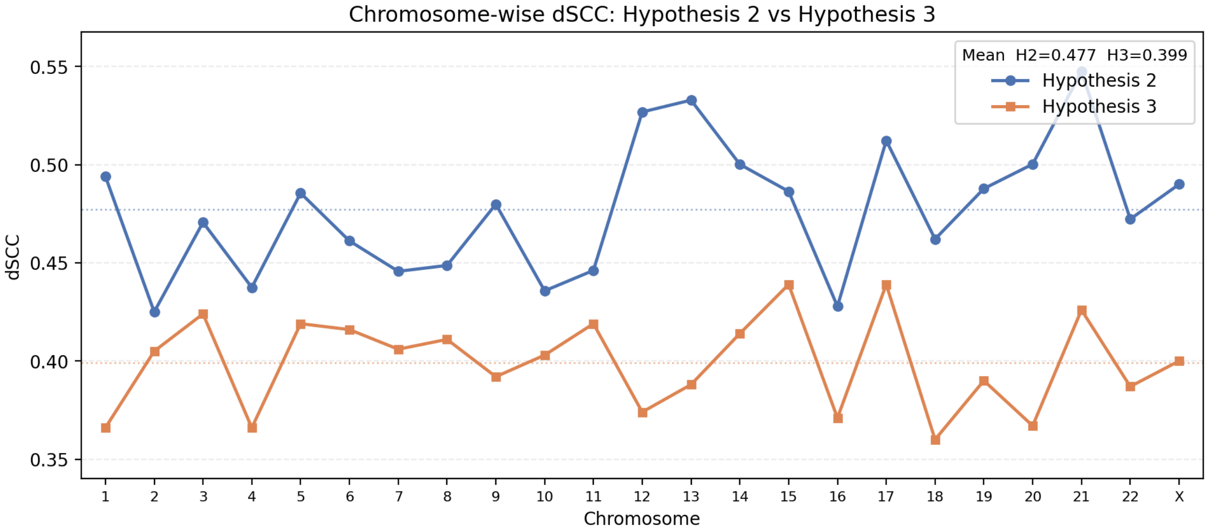


Figure S1: Comparison of chromosome-wise dSCC scores for HiC-LEGO reconstructions generated using domain sets selected by Hypothesis 2 and Hypothesis 3 across all 23 GM12878 chromosomes.

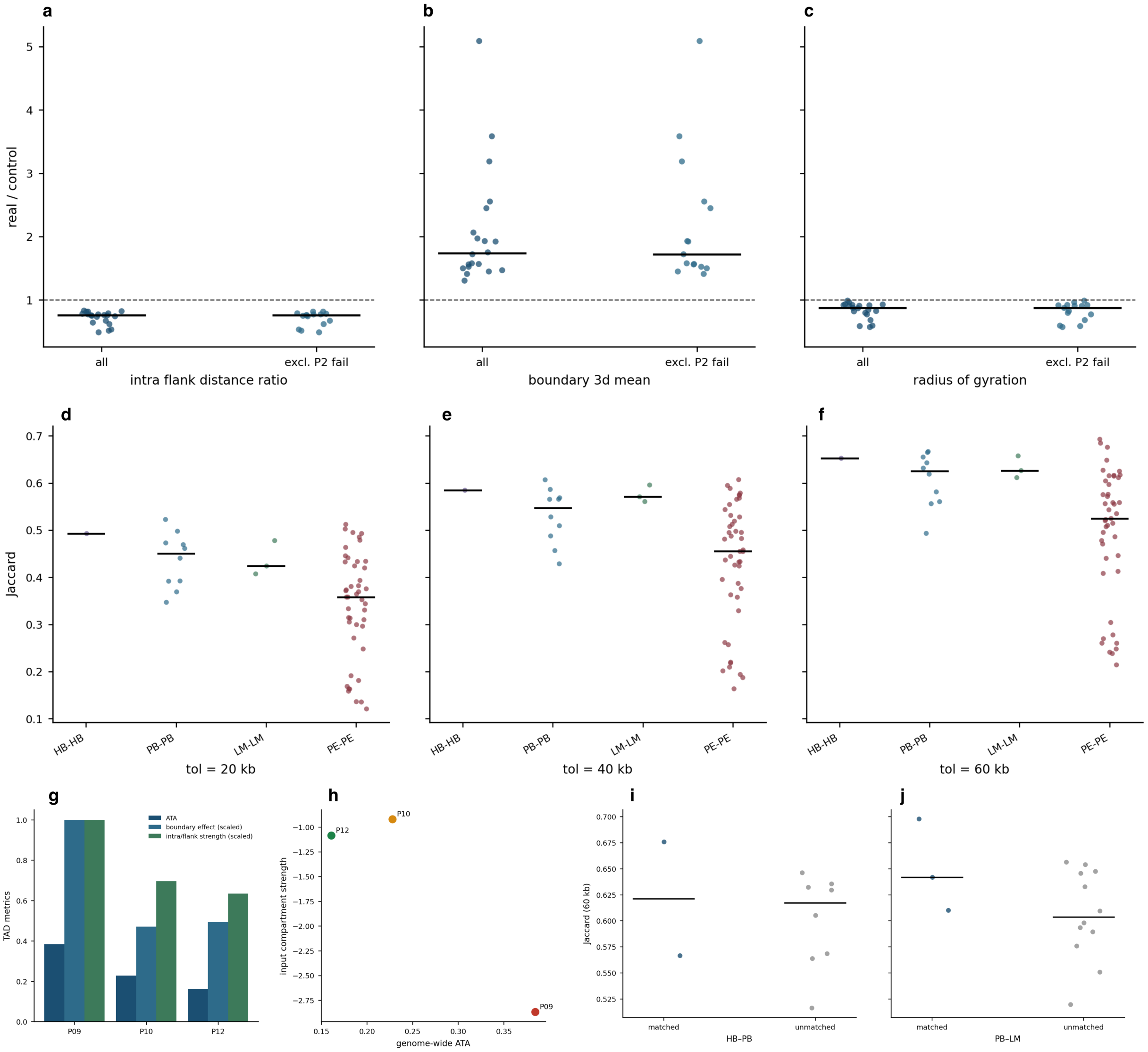


Figure S2: **Sensitivity and supporting analyses for chr13 20-kb breast-cancer validation.** **(a–c)** Sensitivity of supplied-TAD preservation metrics to Phase 2 reconstruction failures. Per-sample median real/control ratios for **(a)** intra/flank 3D-distance ratio, **(b)** direct 3D boundary score, and **(c)** radius of gyration, shown for the full cohort ("all") and after excluding samples that did not outperform the genomic-distance-matched null ("excl. P2 fail"). Horizontal bars indicate group medians; the dashed line indicates a real/control ratio of 1. **(d–f)** Within-stage TAD-boundary Jaccard similarity using matching tolerances of 20 kb **(d)**, 40 kb **(e)**, and 60 kb **(f)**. Points represent pairwise Jaccard scores for HB–HB, PB–PB, LM–LM, and PE–PE comparisons; horizontal bars indicate group medians. **(g,h)** Comparison of TAD and compartment organization among PE samples P09, P10, and P12. **(g)** Genome-wide ATA strength together with scaled regional TAD metrics representing boundary effect and intra/flank strength. **(h)** Input compartment strength plotted against genome-wide ATA strength for the same three samples. **(i,j)** Matched versus unmatched TAD-boundary similarity at a 60-kb matching tolerance for **(i)** HB–PB and **(j)** PB–LM comparisons. Matched pairs represent patient-matched stage comparisons, whereas unmatched pairs represent cross-patient comparisons for the same stage combination. Horizontal bars indicate group medians.

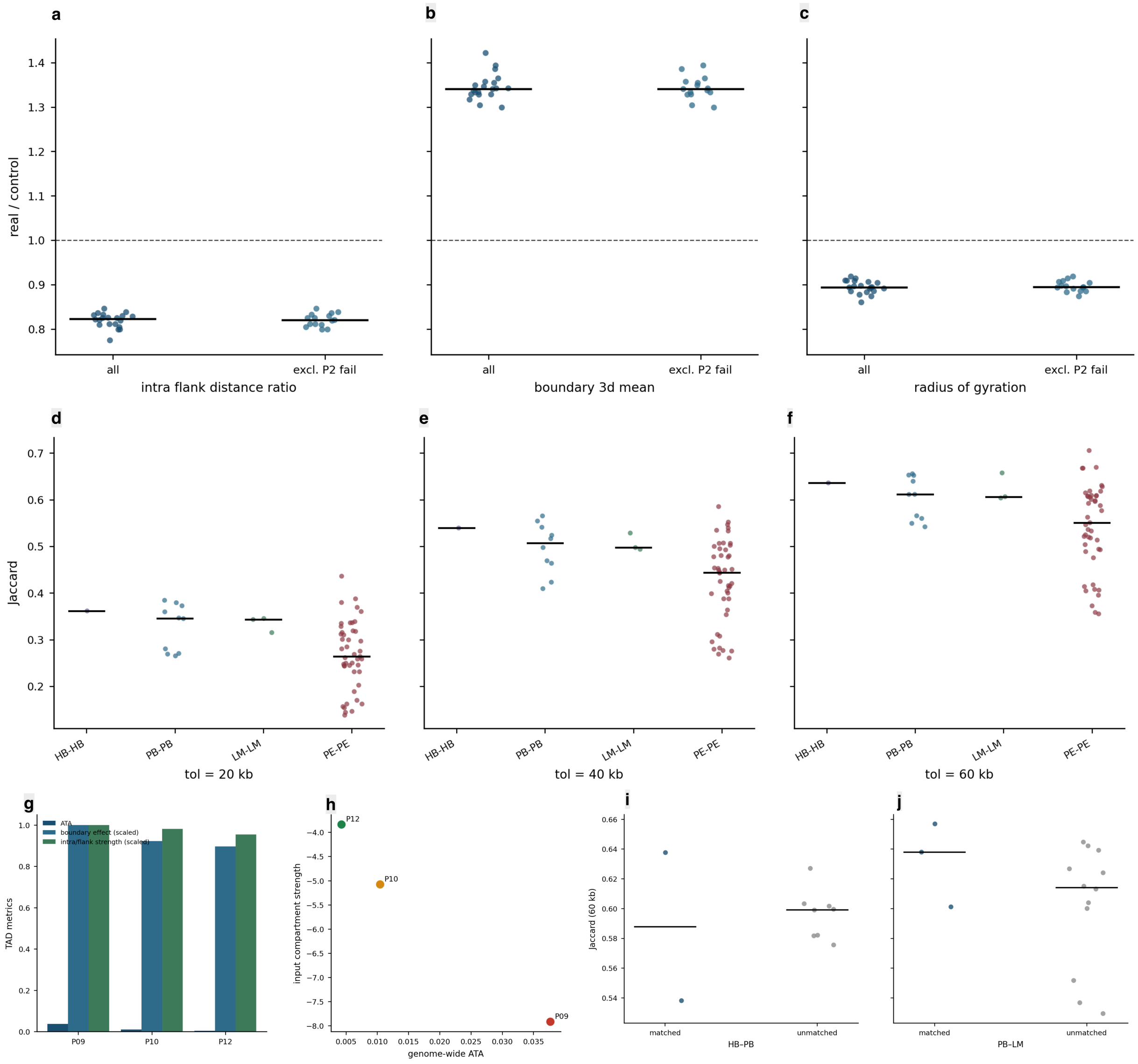


Figure S3: **Sensitivity and supporting analyses for chr13 5-kb breast-cancer validation.** **(a–c)** Sensitivity of supplied-TAD preservation metrics to Phase 2 reconstruction failures. Per-sample median real/control ratios for **(a)** intra/flank 3D-distance ratio, **(b)** direct 3D boundary score, and **(c)** radius of gyration, shown for the full cohort ("all") and after excluding samples that did not outperform the genomic-distance-matched null ("excl. P2 fail"). Horizontal bars indicate group medians; the dashed line indicates a real/control ratio of 1. **(d–f)** Within-stage TAD-boundary Jaccard similarity using matching tolerances of 20 kb **(d)**, 40 kb **(e)**, and 60 kb **(f)**. Points represent pairwise Jaccard scores for HB–HB, PB–PB, LM–LM, and PE–PE comparisons; horizontal bars indicate group medians. **(g,h)** Comparison of TAD and compartment organization among PE samples P09, P10, and P12. **(g)** Genome-wide ATA strength together with scaled regional TAD metrics representing boundary effect and intra/flank strength. **(h)** Input compartment strength plotted against genome-wide ATA strength for the same three samples. **(i,j)** Matched versus unmatched TAD-boundary similarity at a 60-kb matching tolerance for **(i)** HB–PB and **(j)** PB–LM comparisons. Matched pairs represent patient-matched stage comparisons, whereas unmatched pairs represent cross-patient comparisons for the same stage combination. Horizontal bars indicate group medians.

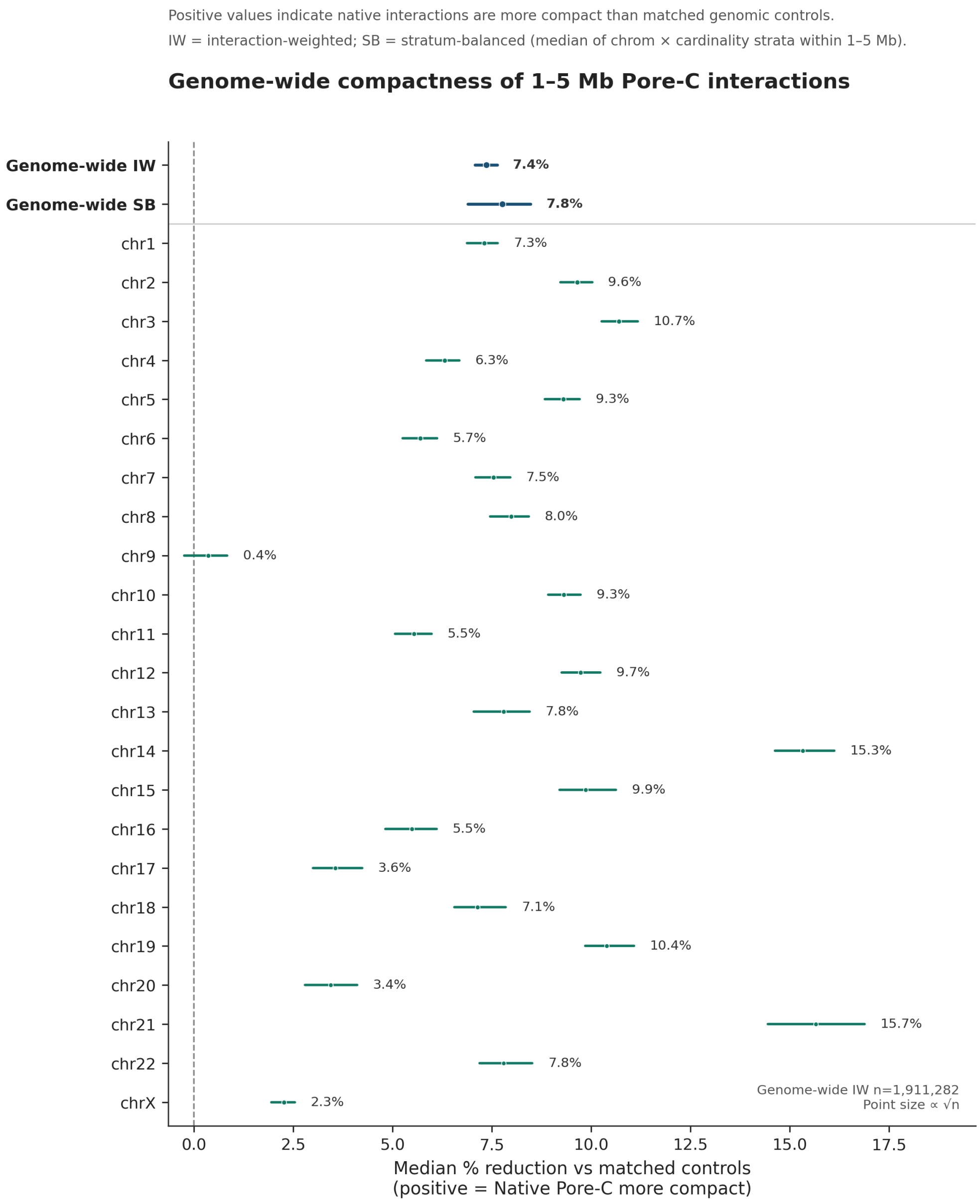


Figure S4: **Chromosome-level compactness of 1–5 Mb Pore-C interactions.** Median percent reduction in maximum pairwise 3D distance between native Pore-C concatemers and matched genomic controls for each chromosome in the 1–5 Mb span stratum. Positive values indicate that native interactions are more compact than their matched controls. Points show median percent reduction and horizontal intervals show 95% confidence intervals. Genome-wide interaction-weighted (IW) and stratum-balanced (SB) estimates are shown at the top. IW gives equal weight to individual concatemers, whereas SB summarizes chromosome-by-interaction-order strata before genome-wide aggregation. The interaction-weighted genome-wide reduction was 7.4%, and all 23 chromosomes showed a positive median reduction.

### 3 Supplementary Tables

Table S1: **Genomic positions and reconstructed 3D distances of GM12878 FISH probes.** Genomic intervals correspond to the L1, L2, and L3 FISH probes used for validation. Euclidean 3D distances between the mapped L1–L2 and L2–L3 probe pairs were calculated from the corresponding HiC-LEGO chromosome structures. Distances are reported in reconstruction model units.

| Chr. | L1 (Mb) | L2 (Mb) | L3 (Mb) | L1–L2 | L2–L3 |
|---|---|---|---|---|---|
| chr11 | 130.72–130.75 | 130.29–130.32 | 129.86–129.89 | 28.71 | 45.98 |
| chr13 | 86.37–86.40 | 85.46–85.49 | 84.55–84.58 | 0.90 | 50.99 |
| chr14 | 71.60–71.63 | 72.20–72.23 | 72.80–72.83 | 23.21 | 53.08 |
| chr17 | 66.76–66.79 | 67.22–67.25 | 67.68–67.71 | 45.33 | 161.06 |

Table S2: **CTCF/RAD21 loop filtering and representative loop used for Fig. 1j–k.** The filtering summary shows the number of loops retained at each step of the CTCF/RAD21 validation. The representative chr22 loop corresponds to the structure shown in Fig. 1k. Reconstructed distances are reported in model units.

| Filtering step | Loops retained |
|---|---|
| Intra-chromosomal GM12878 HiCCUPS loops | 9,448 |
| After blacklist/gap filtering | 9,321 |
| CTCF present at both anchors | 4,990 |
| RAD21 present at both anchors | 4,645 |
| Mapped to distinct 5-kb reconstructed beads | 4,644 |
| Autosomal loops used in Fig. 1j | **4,514** |

| Chr. | Anchor 1 | Anchor 2 | Span | 3D distance | Control median |
|---|---|---|---|---|---|
| chr22 | 23.880–23.890 Mb | 24.100–24.110 Mb | 220 kb | 1.517 | 57.76 |

Table S3: **Breast cancer samples used for chromosome 13 validation.** Samples were analyzed at 20-kb resolution and grouped according to healthy breast (HB), primary breast cancer (PB), liver metastasis (LM), and malignant pleural effusion (PE) stages.

| Sample | Patient | Stage |
|---|---|---|
| GSM8441318_P01_HB | P01 | HB |
| GSM8441321_P02_HB | P02 | HB |
| GSM8441320_P01_PB | P01 | PB |
| GSM8441322_P02_PB | P02 | PB |
| GSM8441324_P03_PB | P03 | PB |
| GSM8441326_P04_PB | P04 | PB |
| GSM8441327_P05_PB | P05 | PB |
| GSM8441319_P01_LM | P01 | LM |
| GSM8441323_P03_LM | P03 | LM |
| GSM8441325_P04_LM | P04 | LM |
| *PE samples P06–P15 continue below* | | |
| GSM8441328_P06_PE | P06 | PE |
| GSM8441329_P07_PE | P07 | PE |
| GSM8441330_P08_PE | P08 | PE |
| GSM8441331_P09_PE | P09 | PE |
| GSM8441332_P10_PE | P10 | PE |
| GSM8441333_P11_PE | P11 | PE |
| GSM8441334_P12_PE | P12 | PE |
| GSM8441335_P13_PE | P13 | PE |
| GSM8441336_P14_PE | P14 | PE |
| GSM8441337_P15_PE | P15 | PE |
